# HHO5: A key orchestrator of dose-dependent nitrogen signaling pathways in Arabidopsis

**DOI:** 10.1101/2025.07.31.667803

**Authors:** Will E. Hinckley, Joseph Swift, Francisco Romei, Jorge P. Muschietti, Shao Shan Carol Huang, Gloria M. Coruzzi, Mariana Obertello

**Affiliations:** Center for Genomics and Systems Biology, Department of Biology, New York University, New York, NY 10003, USA; Plant Biology Laboratory, The Salk Institute for Biological Studies, La Jolla, CA 92103, USA; Instituto de Investigaciones en Ingeniería Genética y Biología Molecular, Dr. Héctor Torres (INGEBI-CONICET), Vuelta de Obligado 2490, C1428ADN, Buenos Aires, Argentina; Departamento de Biodiversidad y Biología Experimental, Facultad de Ciencias Exactas y Naturales, Universidad de Buenos Aires, Int. Güiraldes 2160, Ciudad Universitaria, Pabellón II, C1428EGA Buenos Aires, Argentina; Departamento de Fisiología y Biología Molecular, Facultad de Ciencias Exactas y Naturales, Universidad de Buenos Aires, Int. Güiraldes 2160, Ciudad Universitaria, Pabellón II, C1428EGA Buenos Aires, Argentina

## Abstract

A major goal in agriculture is to engineer crops that can maintain yield with less nitrogen (N) fertilizer input. Major orchestrators of plant responses to N include members of the *HRS1 HOMOLOG* (*HHO*) family of transcription factors (TFs). However, *HHO* TFs have been difficult targets for functional studies *in planta* due to their redundancy. Here, we highlight a unique role for a phylogenetically diverged *HHO* TF, *HHO5,* whose expression is regulated in an N-dose dependent fashion and is specifically expressed in phloem. We found that an *HHO5* single mutant displays significant misregulation of N-dose dependent genes and plant growth rates. HHO5 is also unique as it displays a dual activator/repressor activity on N-dose dependent gene regulation. HHO5 specifically acts as a direct gene repressor when binding DNA targets. In contrast, genes activated by HHO5 include indirect targets regulated by TFs downstream of HHO5 (TF2s). To validate the influence of *HHO5* via its direct TF2s, we used validated TF2 data to build a gene regulatory network that links HHO5-TF2 targets to ∼70% of the N-dose genes regulated by HHO5 *in planta*. By these means, we define HHO5 as a novel dual activator/repressor of plant N-dose signaling.

**Graphical Abstract:** 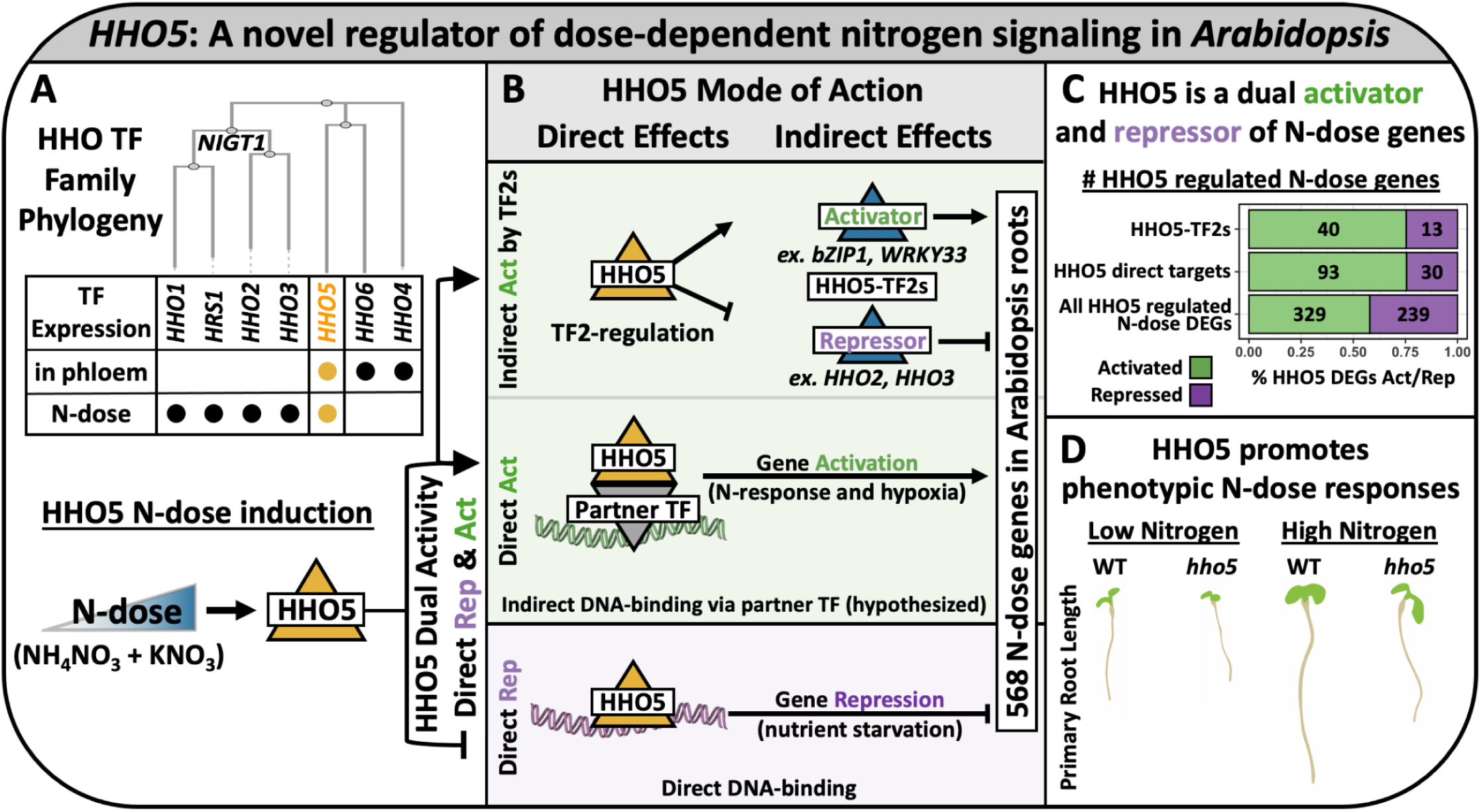

## Introduction

Nitrogen (N) is an essential nutrient for plant growth. Biomass, grain yield, root architecture, and photosynthesis are all impacted by N-availability. Furthermore, while the synthesis of nitrate through the Haber process ushered in the Green Revolution, the production of N-fertilizers is costly and holds negative environmental consequences. For these reasons, the demand for crops that require less N-fertilizer for growth is rising^1^.

N-adaptive responses are initiated at the molecular level, where environmental stimuli trigger receptor mediated signaling cascades, inciting changes in gene regulation, which in turn alters the course of plant development. While the genes that make up such signal transduction pathways are under intense scrutiny, how signal transduction pathways relay messages *quantitatively* receives less attention. N-signals are highly dynamic, varying across the dimensions of both dose and time. Our previous studies integrated dose and time treatments to reveal that temporal N-dose transcriptome signals can be modelled by simple Michaelis Menten kinetics. Increasing N-dose leads to a proportional change in the rate of gene expression activity, transcriptome-wide^2^. This dose-dependent transcriptional response is driven by transcription factor (TF) activity. We validated that perturbations of the TF *TGA1* led to proportional changes in the rate of N-dose response *in planta,* supporting its role in modulating the transcriptional kinetics of N signaling.

An advantage of studying plant responses to N-dose over time is the uncovering of early acting transcriptional regulators. The first TFs to respond to N-treatments (within minutes) are assumed to play key roles in stimulating downstream metabolic pathways. Key early responding TFs to N signals include *CRF4* and several members of the *NIGT1/HRS1*(*HHO*) family, such as *HHO2, HHO3,* and *HHO5*^3^. These TFs are important candidate regulators of N-signaling, as their genetic perturbation could enhance the plasticity and efficiency of *in planta* N-dose responses.

The *HHO (HRS1-HOMOLOG)* family is a subset of the G2-Like class of plant MYB TFs, known for its role in repressing N responses^4^. Of the seven HHO genes, four TFs (*HRS1, HHO1*, *HHO2, HHO3*) constitute a group of closely related TFs called the *NITRATE-INDUCIBLE, GARP-TYPE TRANSCRIPTIONAL REPRESSOR1* proteins (*NIGT1s*)^5^.

*NIGT1* TFs influence N-signals by redundantly repressing high-affinity nitrate transporters^6^. *NIGT1* TFs only influence N-related plant phenotypes when mutated in higher-order combinations due to their redundancy. It could be agriculturally important that these *NIGT1* TFs influence N-related plant growth and development, but their redundancy makes them difficult targets for gene editing applications.

In contrast, *HHO4*, *HHO5*, and *HHO6* are more phylogenetically diverged from the *NIGT1* TFs, and their role in N-signaling has not been studied. Notably *HHO5* was previously shown to act as a gene repressor in the floral meristem^7^. In our previous studies, *HHO5* emerged as one of the earliest transcriptional responders to N-treatments (within 10 minutes)^2,3^. Moreover, *HHO5* stands out, as we previously identified *HHO5* to have a highly conserved N response between rice and Arabidopsis^8^. *HHO5* is thus a strong candidate regulator of plant N-dose signaling.

In this study, we characterize the dual activating and repressing role of *HHO5* in regulating N-dose responses *in planta*. We found that HHO5 regulates the N-dose response of hundreds of genes *in planta*. While the *HHO NIGT1* repressors require additive mutations to influence plant growth, single mutation of the more phylogenetically diverged *HHO5* decreased plant growth in a N-dose dependent manner. This *hho5* mutant growth-activating phenotype contrasts the known repressive role of HHO TFs. To better understand this phenomenon, we characterized the HHO5 regulatory mode of action. We found that HHO5, like the *NIGT1* TFs, acts as a direct gene repressor when binding DNA. However, HHO5 also activates many N-dose genes (transiently in cells, and *in planta)*. Therefore, HHO5 activation of N-dose response*s* must be the consequence of indirect regulatory effects, such as interacting partner TFs and/or downstream TFs (TF2s).

To validate the influence of HHO5 indirect activity, we built a gene regulatory network (GRN) using a network walking approach. Network walking allowed us to map the indirect effects of HHO5 by incorporating the regulatory activity of secondary TF2s downstream of HHO5. The combined direct and indirect effects of *HHO5* explained the *in planta* regulation of ∼70% of the HHO5 targeted N-dose genes. Thus the dual activator/repressor transcriptional activity of HHO5 results in unique plant N-dose responses that are not associated with other HHO TFs. We also found that *HHO5* is expressed specifically in phloem cells, where the *NIGT1* repressors are not expressed. Therefore, the spatial divergence of *HHO5* may further explain why *hho5* mutants display unique phenotypic responses to N-dose. The finding that HHO5 activates N-dose responses *in planta* contrasts the canonical repressive role of other HHO TFs, and highlights the importance of studying a more phylogenetically diverged N-regulating TF.

## Results

To investigate how HHO5 regulates Arabidopsis responses to changing N-dose, we grew plants on N-depleted media and transiently treated WT (Col-0) and two *hho5* T-DNA mutants *(hho5.1, hho5.2)* for 2 hours with each of six N-doses, ranging from 0 mM to 60 mM N (supplied as KNO_3_ + NH_4_NO_3_). The highest N-dose reflected the amount of N present in standard MS media^9^. We included a 20 mM KCl treatment to control for the effects of potassium in KNO_3_ (**Supp Figure 1**). Transient treatment with N ensured we captured genes involved in early N sensing, which was important given HHO5 is one of the earliest TFs to respond to N-stimulus^3^. Whole root transcriptomes were then analysed with RNA-seq.

We first identified transcriptome-wide N-dose responses in WT. After quality control filtering steps (see Materials and Methods) we found **4,464** genes whose expression was N-dose responsive relative to the KCl treatment control. **1,702** genes were upregulated, and **2,272** genes were downregulated as N-dose increased (**Figure 1A** and **Table S1**). Some genes were activated or repressed by N-dose, but also separately activated or repressed specifically in response to N-starvation (0 mM N, **Supp Figure 2**). Genes activated by N-dose were enriched for nitrate assimilation, hypoxia, and response to cytokinin. This is consistent with previous literature showing positive associations between N and cytokinin signals^10^. Hypoxia signals, via reactive oxygen species (ROS), have been shown to be important for *NIGT1* TFs in regulating N-signaling^6^. Alternatively, genes repressed by N-dose were enriched for defense related terms, calcium transport, and water responses. These terms are also relevant to N-signaling (**Figure 1B** and **Table S2**)^11,12,13^.

**Figure 1:**
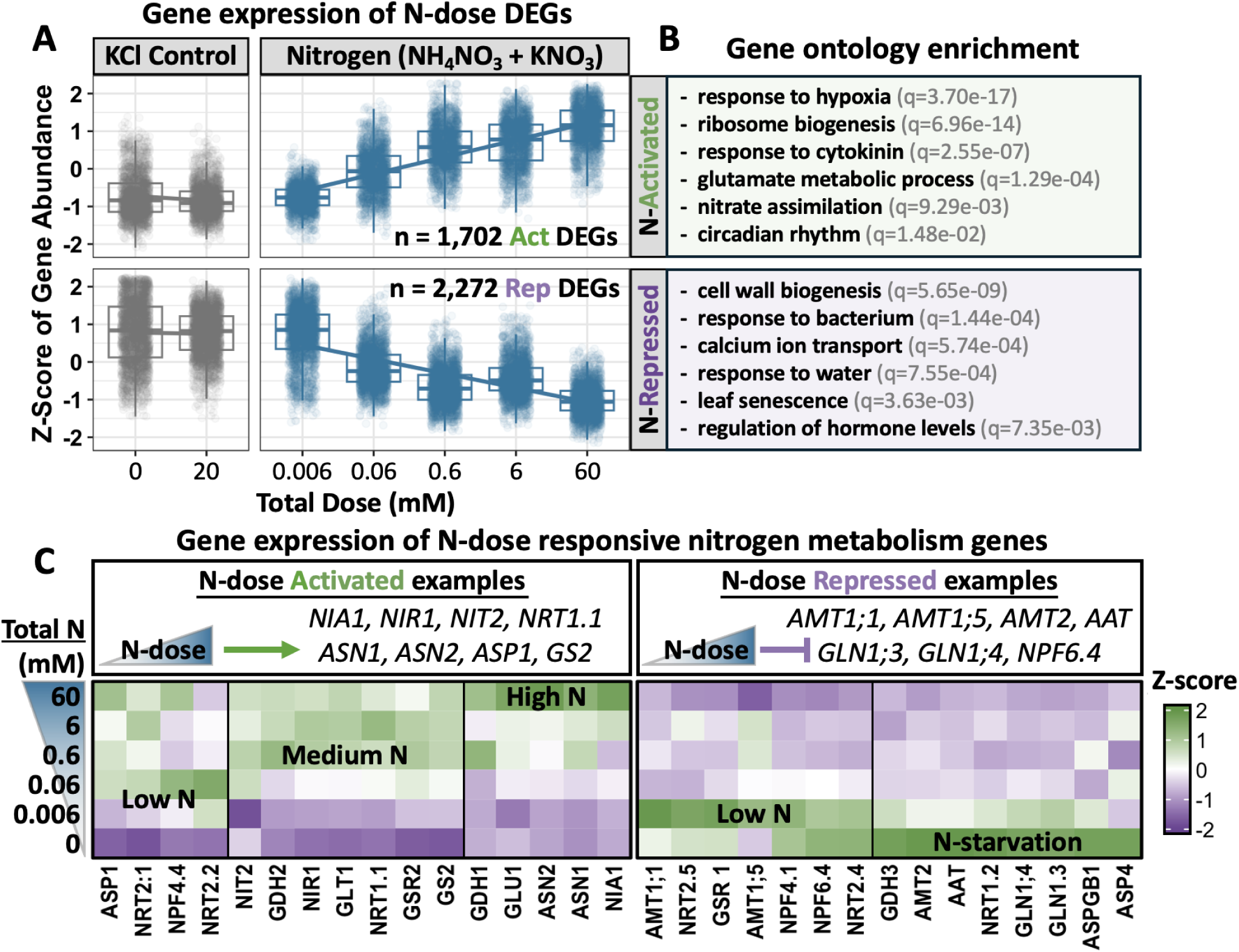
The *Arabidopsis* root transcriptome is sensitive to N-dose signals. A) Z-score scaled expression of genes activated (on top) or repressed (on bottom) by N-dose signals relative to KCl treatment controls. There are 4,464 N-dose DEGs total (DESeq2 p.adj < 0.05). Total mM N-dose corresponds to 1xNH_4_ (<u>NH_4_</u>NO_3_) and 2xNO_3_ (K<u>NO_3_</u> and NH_4_<u>NO_3_</u>). For example, 60 mM Total N reflects treatment with 20 mM NH_4_NO_3_ and 20mM KNO_3_. B) Top non-redundant enriched GO terms in the N-dose activated or repressed gene clusters, next to the q-value of enrichment significance. C) Z-score scaled expression of known N metabolism genes across N-doses, ordered by hierarchical clustering.

As many N-metabolism-related GO Terms were enriched in the N-dose differentially expressed genes (DEGs), we next queried the expression of known N-metabolism genes across N-doses treatments. Many key genes involved in the transport, reduction, and assimilation of N were N-dose responsive (**Figure 1C**). Over half of the 59 KEGG N-metabolism pathway genes (52.5%) genes displayed significant N-dose responses (31/59, p-value = 2.90e^-07^)^14^. For example, genes related to nitrate uptake (*NRT1*.*1*, *NRT2*.*1*), N-reduction (*NIA1*) and N-assimilation (*ASN1*, *ASN2, GLT1*, *GDH1* and *GS2*) were N-dose upregulated. Genes related to N starvation, such as *NRT2.4* and *GDH3*, and ammonium transport (*AMT1.1* and *AMT1.5*) were downregulated in response to N-dose, agreeing with previous studies of N-responses^5^. We were also able to distinguish N-metabolism genes activated specifically by N starvation (0 mM N) from genes activated more strongly by extremely low N supply (0.006 mM N, **Figure 1C**). Together, these results show that canonical N-metabolism genes in the Arabidopsis root transcriptome are sensitive to N supply as a function of dose.

The N-responsiveness of some HHO TF family members has been previously studied, especially for the *NIGT1* TFs. Here, we analyzed the N-dose response of the entire Arabidopsis HHO family. We found that five of the seven HHO TFs significantly responded to N-dose (all but *HHO4* and *HHO6*, **Figure 2A**). Among the non-*NIGT1* HHO TFs, *HHO5* is the only TF that exhibits a significant transcriptional response to N-dose in our dataset. To investigate the spatial expression pattern of HHO5 in roots, we queried the gene expression of the HHO TF family across an Arabidopsis bulk RNA-seq root cell-type atlas^15^. Interestingly, the clustering of gene expression profiles across root cell types mimicked the HHO family protein phylogeny (**Figure 2B**). While the *NIGT1* TFs were expressed broadly across many cell types, including cortex, endodermis, and hair cells, the non-*NIGT1* TFs (*HHO4, HHO5*, and *HHO6*) displayed a more specific expression pattern, predominantly in the phloem and companion cells.

**Figure 2:**
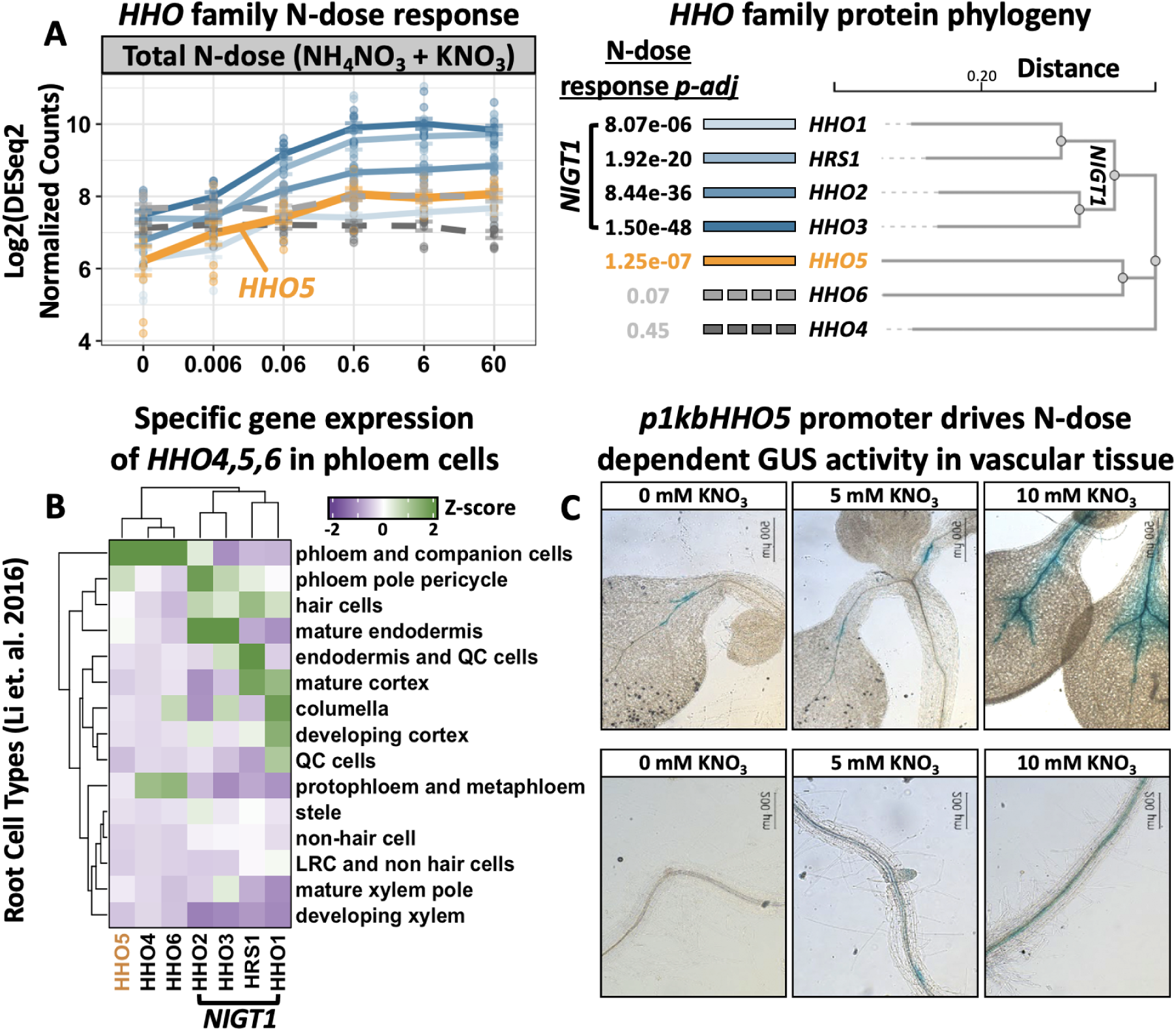
HHO5 is expressed in phloem and responds to N-dose. A) Log2 DESeq2-normalized gene expression of the 7 HHO Family TFs across N-doses. Lines are colored by annotation in the protein phylogeny, showing the adjusted p-value for N-dose response from DESeq2. B) Z-score scaled gene expression of the HHO TFs across the bulk RNA-seq root cell type atlas from Li et al 2016^15^. C) N enhances activity of p1kbHHO5::GUS expression of 10 days old shoots and roots across three N conditions.

Notably, *HHO5* clustered away from *HHO4* and *HHO6*, as *HHO5* is not expressed in the protophloem and metaphloem. *HHO5* instead had specific gene expression in the phloem pole pericycle where *HHO4* and *HHO6* are not expressed (**Figure 2B**). Taken together, these findings indicate that HHO5 diverges from the *NIGT1* TFs not only in its protein sequence but also with its distinct spatial expression in phloem cells. To validate the localization of *HHO5*, we completed GUS reporter activity experiments using the 1 kb upstream promoter of *HHO5* (*p1kbHHO5*). *p1kbHHO5* driven GUS activity increased in response to higher N-doses in both shoots and roots, and was localized to the vascular tissues (**Figure 2C**). In summary, *HHO5* gene expression significantly responds to N-dose signals, and is preferentially localized to phloem cells.

### HHO5 regulates genome-wide N-dose signals

Our next goal was to evaluate the role of HHO5 in regulating *in planta* N-dose signals. We found that mutating *HHO5* resulted in mis-regulation of hundreds of N-dose genes *in planta*. We first used transcriptome data from *hho5* T-DNA mutants to identify genes that are differentially expressed (DEGs) relative to the WT control^16^. We also confirmed *HHO5* gene expression response was significantly perturbed in the *hho5.1* and *hho5.2* mutants (**Supp Fig 3A**). 3,605 and 4,338 genes were differentially expressed in the *hho5.1* and *hho5.2* mutants, respectively. 2,894 DEGs were shared between the two mutant backgrounds, representing a highly significant DEG overlap between the two mutant alleles (**Figure 3A, Supp Fig 3B, Table S3**). To ensure we found high confidence HHO5 DEGs, we only considered a gene to be regulated by HHO5 *in planta* if it was significantly perturbed in both mutant alleles. Furthermore, the DEG log2 fold change (mutant/WT) for these shared DEGs were highly correlated between the two *hho5* mutant alleles (R2 = 0.989, p-value < 2.2e^-16^, **Supp Fig 3C**). Therefore, we robustly identified thousands of genes regulated either directly or indirectly by HHO5 *in planta*.

**Figure 3:**
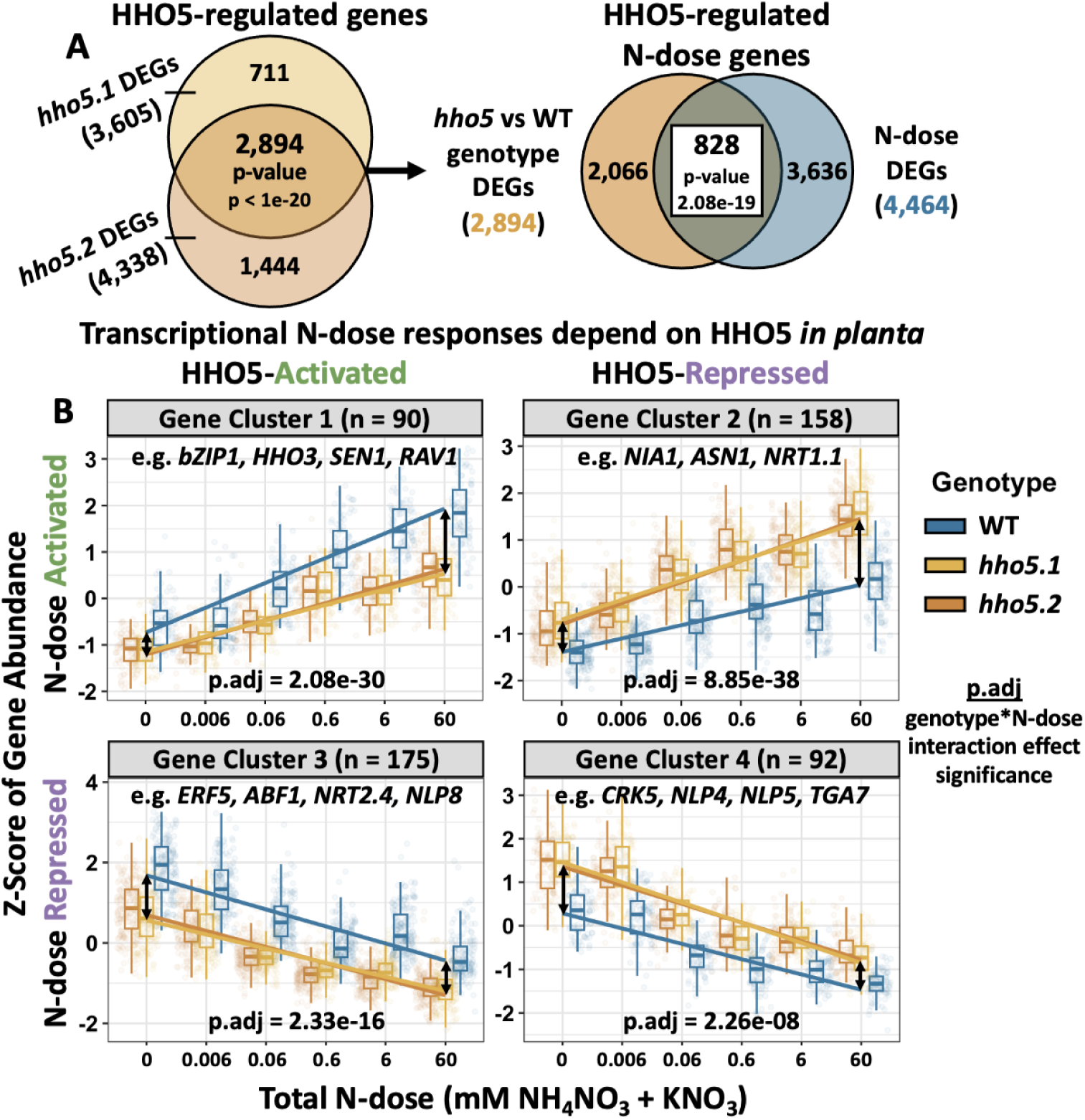
HHO5 regulates the N-dose response of hundreds of genes *in planta*. A) Intersection of DEGs from two hho5 T-DNA mutants (DESeq2 p.adj < 0.05) for the genotype effect. Intersection significance is the result of a hypergeometric test. 2,894 genes are regulated in both mutants, 828 of which intersect with the 4,464 N-dose DEGs from Fig 1. B) Expression profiles of 4 largest clusters of HHO5 N-dose DEGs.

Importantly, a significant portion of the HHO5 *in planta* DEGs also respond to N-dose treatments (828 genes, p = 2.08e^-19^), demonstrating that HHO5 directly or indirectly regulates a substantial fraction (∼19%) of the N-dose responsive transcriptome (**Figure 3A, Table S4**). To visualize the HHO5-regulated N-dose DEGs, we clustered and plotted the expression of the DEGs across genotypes and N-doses. Most genes fell into four clusters which were either activated or repressed by HHO5 and/or N (**Figure 3B**). For example, genes in Cluster 1 increase gene expression with increasing N-dose (blue line). Cluster 1 genes were expressed lower in *hho5* mutants (orange lines) relative to WT (blue line). Therefore Cluster 1 genes, such as the N-signal activating hub TF *bZIP1*, are activated both by HHO5 and by N-dose *in planta*.^17^ In contrast, genes in Cluster 2 are activated by N-dose but are repressed targets of HHO5.

Cluster 2 included important N-metabolism genes *NRT1.1, NIA1*, and *ASN1*. HHO5 repression of N-metabolism genes *in planta* is consistent with previous literature stating HHO TFs act as N-signal repressors^6^. However, while HHO TFs are described as repressors, HHO5 also activated the expression of many N-dose genes (**Clusters 1 and 3, Fig 3B**). This analysis determined that HHO5 can both activate and repress N-dose genes *in planta*.

To determine if transcriptional N-dose responses were significantly perturbed by *HHO5* mutation, we used an ANOVA to test for a genotype*N-dose interaction effect within these gene clusters. The ANOVA revealed in all 4 clusters that the N-dose response of these genes, on average, significantly depends on the level of *HHO5*. The genotype*N-dose interaction effect seemed slightly stronger amongst genes activated by N-dose compared to the N-dose repressed genes **(Figure 3B).** Therefore, HHO5 can both activate and repress the N-dose response of hundreds of genes *in planta*.

### HHO5 promotes phenotypic growth responses to N-dose signals

To further validate the role HHO5 plays in regulating N-dose signals *in planta*, we investigated the impact HHO5 had on the ability of *Arabidopsis* to grow in response to different N-doses. We grew wild-type seedlings (Col-0) and both *hho5.1* and *hho5.2 null* T-DNA mutant lines on 4 different N-doses that ranged from a growth limiting dose of 0.1 mM to a moderate dose of 10 mM KNO_3_ (see Materials and Methods). As expected, we found wild-type plant phenotypes were plastic in response to varying N availability, with both plant dry weight and root length adapting to the levels of N provided (**Figure 4**).

**Figure 4:**
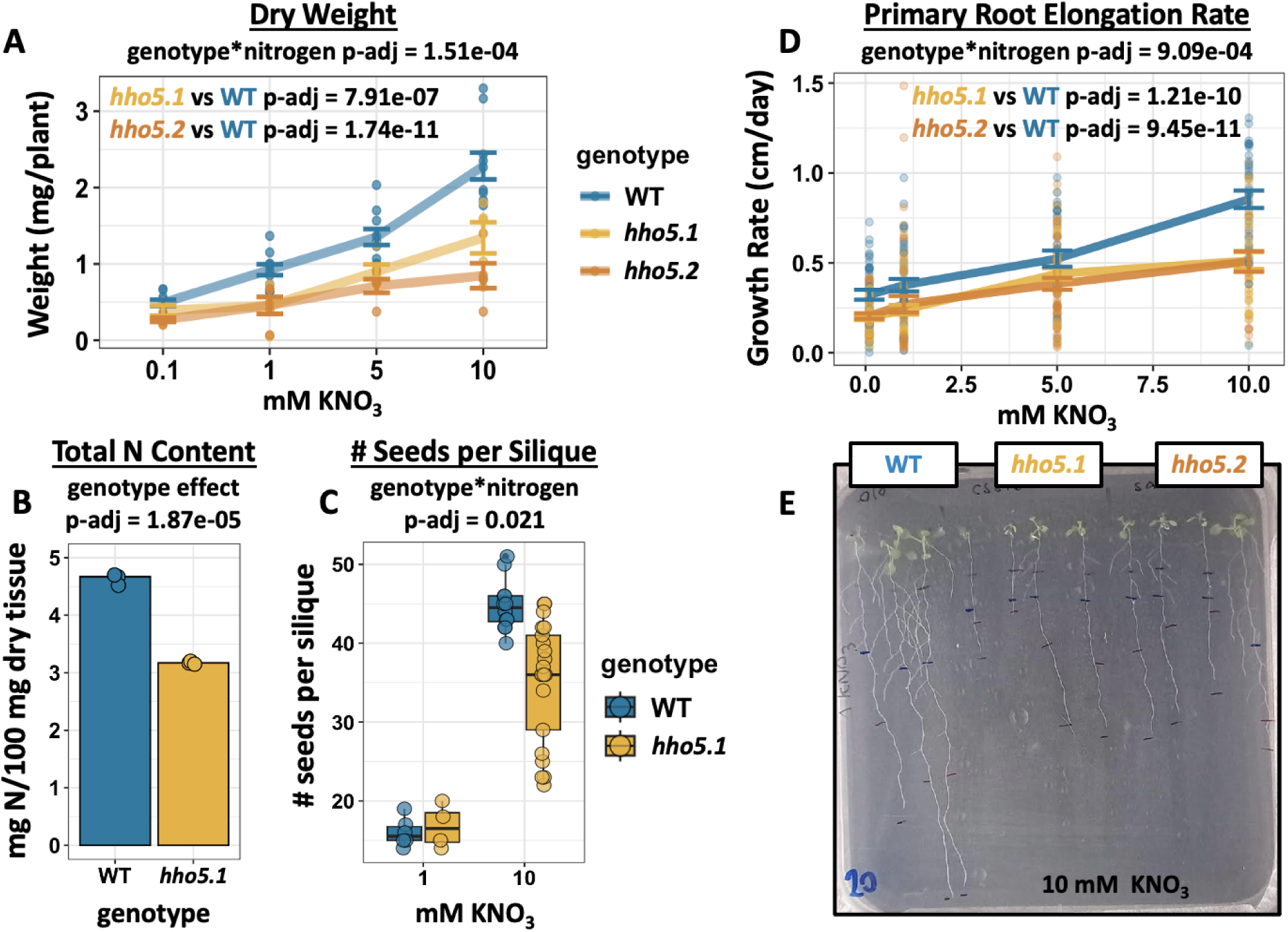
HHO5 promotes phenotypic N-dose responses. A) Plant dry weight across N-doses in WT and hho5 mutants, where an ANOVA was performed to test for a significant genotype*N-dose interaction effect. Plant fitness is worsened in the absence of HHO5, where B) total N content in seeds, and C) the number of seeds per silique, are decreased in the hho5.1 mutant compared to WT plants. D) Primary root growth rate (cm growth/day) vary across N-dose, and are perturbed in both hho5 T-DNA mutants. E) Image of root lengths at 10 mM KNO_3_.

Upon perturbing *HHO5*, we found that the plant’s ability to respond to N-dose was impaired, leading to a significant reduction in shoot biomass (**Figure 4A, Supp Fig 6**).

Therefore, HHO5 activates N-dose responses both transcriptionally, and at the phenotype level. Specifically, under the growth limiting N condition of 0.1 mM, the absence of HHO5 resulted in a 36% reduction in shoot biomass. This effect was further intensified under N-replete conditions (10 mM), leading to a 51% reduction in shoot biomass. A significant genotype*N-dose interaction effect was found for shoot dry weight, indicating the growth of *hho5* mutants is impaired in a N-dose dependent manner (p<1e^-04^). These growth defects resulted in a strong reduction in seed N content, as well as a reduction in the number of seeds per silique when comparing the *hho5.1* mutant against the WT (**Figure 4B-C**). We also confirmed these phenotypes in response to 4 doses of ammonium nitrate to ensure mutant phenotypes were detected in response to different sources of N (**Supp Fig 5**).

We previously found that N-dose responses are dynamic, and that perturbing the TF *TGA1* affects the *rate* of plant growth in response to N^2^. To determine if growth rates in response to N-dose are under the control of HHO5, we measured the primary root length at 4 time points over 3 day periods (**Supp Fig 6**). Indeed, the growth rate of the primary root increases with increasing N-dose (**Figure 4D**). This increase in growth rate is significantly perturbed in the absence of *HHO5*, meaning HHO5 is required for maintaining primary root growth rate plasticity in response to N-dose (**Figure 4E**). In summary, growth defects in both *hho5.1* and *hho5.2* mutants were more intensified with increasing N-dose. *HHO5* is thus important for promoting plant plasticity and fitness in response to varying N-doses. However, since N-dose responses were not entirely abolished in the mutants, there must be additional N-dose response mechanisms that are independent of *HHO5*.

### HHO5 direct DNA-binding is associated with target gene repression

The finding that HHO5 activates phenotypic responses to N-dose was surprising given the canonical repressive role of HHO TFs. Additive mutants of the redundant NIGT1 TFs were previously shown to display the opposite (repressive) phenotype^6^. As *HHO5* is phylogenetically diverged and has a distinct spatial expression profile compared to the repressive *NIGT1* TFs (**Figure 2**), we hypothesized that HHO5 may have a different gene regulatory mode of action.

To mechanistically understand how HHO5 regulates *in planta* N-dose responses, we mined *HHO5* over-expression data from the TARGET TF perturbation assay^3,18^. Briefly, the TARGET assay involves transient over-expression of a TF, in this case *HHO5,* fused to a rat glucocorticoid receptor domain (GR) in Arabidopsis protoplasts. The GR domain of the GR:HHO5 fusion interacts with HSP90, which sequesters the fusion protein into the cytoplasm. Treatment with dexamethasone (DEX) disrupts the GR:HSP90 interaction, allowing for precise control of GR:HHO5 nuclear import. After this, RNA-seq is performed to identify differentially expressed genes in response to nuclear import of GR:HHO5 relative to an empty vector (EV, no TF, GR-only control). Crucially, before nuclear localization, cycloheximide (CHX) is added to prevent translation of downstream, secondary transcripts (such as other TFs) that are regulated by the GR:TF fusion. By these means, the TARGET assay can reveal the genes directly downstream of the GR:TF fusion^18^. TARGET also has an advantage of capturing early target genes that could be missed *in planta*, as the transient assay can be stopped after only hours of TF-nuclear import.

TARGET revealed that HHO5 directly activated **908** genes and directly repressed **615** genes in protoplasts (**Figure 5A, Table S5**), reinforcing our hypothesis that HHO5 is both a transcriptional activator and repressor of N-signaling genes. Genes directly activated by HHO5 were enriched for N related terms, hypoxia, and defense, similar to genes activated by N-dose *in planta* (**Table S6**). In contrast, HHO5 directly repressed genes related to nutrient starvation, which has been seen for other HHO TFs^6^. In total, *HHO5* and N-stimuli regulate pathways such as N-metabolism, hypoxia, defense, and starvation in the same direction.

**Figure 5:**
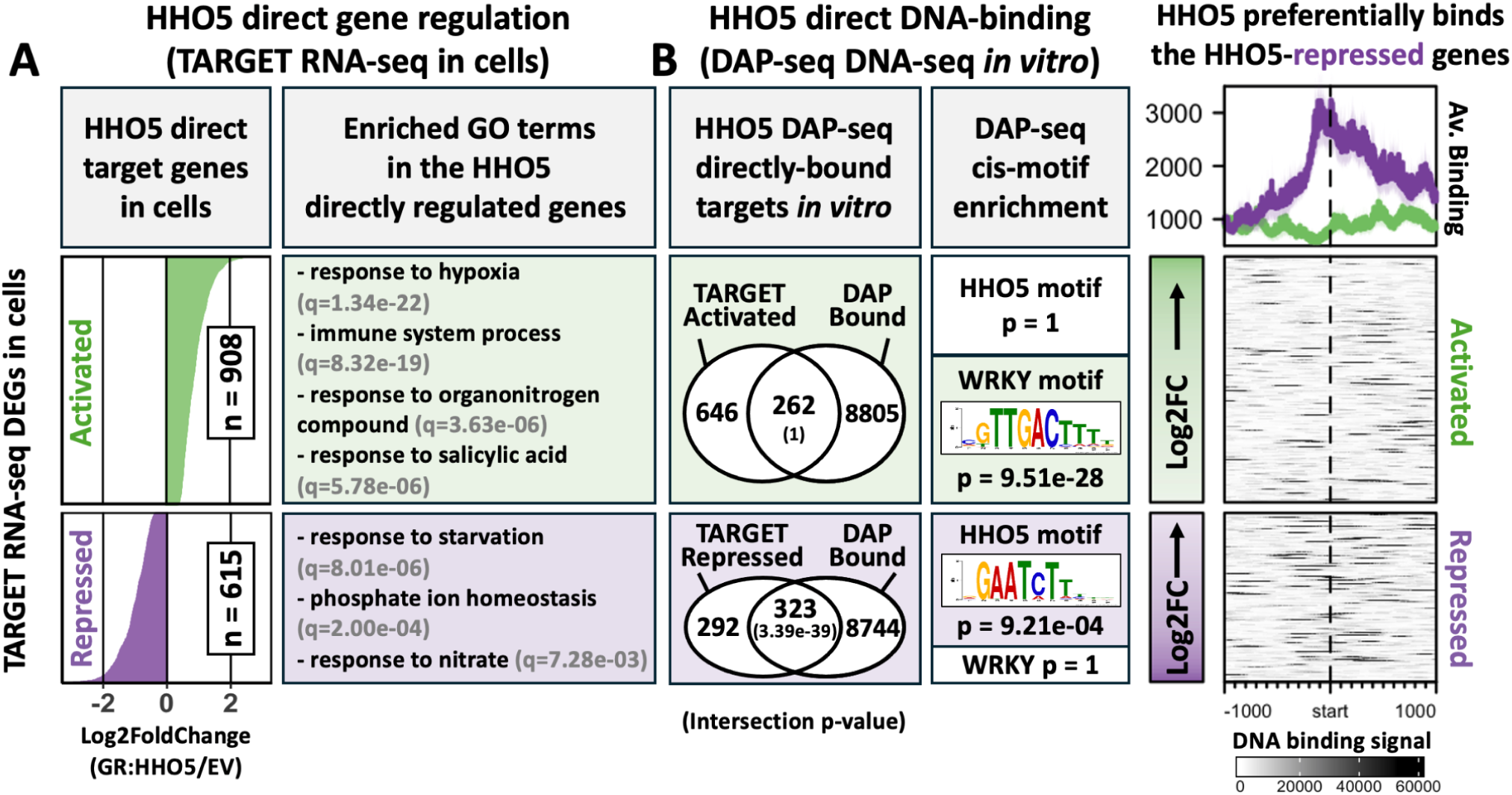
HHO5 direct DNA binding is associated with target gene repression. A) HHO5 direct target genes in cells from the TARGET system^3^ and associated GO Terms, with q-values for enrichment significance. B) HHO5 DAP-seq^19^ direct DNA Binding intersection with TARGET RNA-Seq DEGs, with p-values from a hypergeometric test for significance. Plotted next to HHO5 DAP-seq binding signal over the TARGET regulated genes. The TARGET regulated genes are ordered by Log2 fold change (GR:HHO5/GR:EV), and represent +/- 1kb of sequences around the gene transcription start sites.

We next sought to characterize HHO5 DNA binding using published DAP-seq data^19^.

In DAP-seq, the TF of interest is fused to a HALO tag, and the HALO:TF fusion is expressed *in vitro*. After incubation with a genomic DNA library, affinity-precipitation of the HALO:TF fusion captures genome-wide DNA binding sites. Because no other TFs are expressed *in vitro*, DAP-seq binding signal represents direct DNA binding^20^. We found that hundreds of HHO5 direct target genes from the TARGET assay were also directly bound by HHO5 in DAP-seq (**Figure 5B, Table S7)**. Interestingly, there is a significant intersection between the HHO5 DAP-seq bound genes and the HHO5-repressed genes, while no such enrichment was observed for the HHO5 activated genes. This indicates, similarly to other HHO TFs, that HHO5 direct DNA binding is associated with gene repression^7^.

DAP-seq is an *in vitro* assay, meaning some DNA-binding peaks may not occur *in vivo* due to epigenetic or spatial constraints. To further confirm that HHO5 prefers binding to HHO5 repressed genes, we queried enrichment of the known HHO5 DAP-seq DNA-binding motif among the HHO5 TARGET-regulated genes. The canonical HHO5 DNA-binding site is significantly enriched in the HHO5 repressed genes, but not the HHO5 activated genes (**Figure 5B**). Interestingly, the HHO5 activated genes instead display significant enrichment for the WRKY TF family binding motif. To mitigate thresholding issues from peak calling, we also quantified HHO5 DAP-seq DNA binding over the HHO5 regulated genes. Not only was HHO5 direct DNA binding signal much stronger among the HHO5 repressed genes, but the binding signal was also localized to the target gene transcription start sites (labeled as start, **Figure 5B**). This was also the case for HHO5 ampDAP-seq, in which the DNA library is PCR amplified to remove endogenous DNA methylation. Thus, DNA methylation did not have an obvious effect on HHO5 DNA binding to the HHO5 repressed genes (**Supp Fig 7**).

As an example target gene, we show that HHO5 repressed *NRT1.1* (AT1G12110) both in protoplasts and *in planta*, and HHO5 directly bound the promoter and gene body of *NRT1.1* in DAP-seq (**Supp Fig 8**). All together, these analyses suggest that direct DNA binding by HHO5 is associated with target gene repression. We repeated this analysis for other HHO TFs with available data and found the same relationship between direct DNA binding and HHO5-target gene repression (**Supp Fig 7**). Therefore, the functional divergence of HHO5 from other HHO TFs is not related to its direct DNA binding activity, as directly repressive DNA binding is conserved across the HHO TFs.

It became our goal to better understand gene activation directly downstream of HHO5. Gene activation downstream of HHO5 is likely due to indirect transcriptional effects, such as indirect DNA binding via partner TFs (heterodimerization), or the regulatory effects of secondary TFs downstream of HHO5 (TF2s). While indirect DNA binding via a Partner TF could be an interesting mechanism of HHO5 target-gene activation, TF-DNA binding does not always correlate to gene regulation^21^. Instead, we chose to focus our study on HHO5 TF2s, with which we could robustly validate the indirect regulation of N-dose genes downstream of HHO5.

### A GRN links HHO5 gene regulatory activity in cells to N-dose responses *in planta*

To better understand the regulatory effects of HHO5 on N-dose responses, we intersected HHO5 direct TARGET DEGs in cells with the HHO5 mutant *in planta* DEGs. HHO5 regulates 828 N-dose genes *in planta* (**Figure 3**), and 135 of these 828 genes are also directly regulated by HHO5 in cells (TARGET assay) (**Supp Fig 9**). The 135 direct N-dose targets of HHO5 are enriched for response to N compound (q = 5.92e^-04^), response to hypoxia (q = 6.29e^-07^), and response to starvation (q = 1.34e^-02^). Therefore HHO5 directly regulates N-dose genes related to N-metabolism and nutrient starvation both in cells and *in planta*.

We characterized the indirect regulatory effects of HHO5 on N-dose responses using a network walking approach^22^. We reasoned that genes regulated by HHO5 *in planta*, but not directly regulated by HHO5 in cells, are likely indirect target genes of HHO5. To validate the HHO5 indirect regulation of N-dose genes *in planta* with network walking, we used TARGET gene regulation data for TFs downstream of HHO5 (TF2s). Of the 22 N-dose responsive TF2s directly downstream of HHO5 both in cells and *in planta*, 10 TF2s had TARGET gene regulation data (**Figure 6**) in our ConnecTF database^23^. Three additional TFs downstream of the HHO5 TF2s also had TARGET data (TF3/4s). TARGET-validated direct gene regulation from these 14 TF2/3/4s further explained 418 of the 828 HHO5 *in planta* N-dose DEGs (**Table S8**). In total 553 of the 828 HHO5 regulated N-dose DEGs (66.8%) were validated using TARGET gene regulation data, with a majority of these DEGs being indirect HHO5 target genes under the control of HHO5-TF2s.

**Figure 6:**
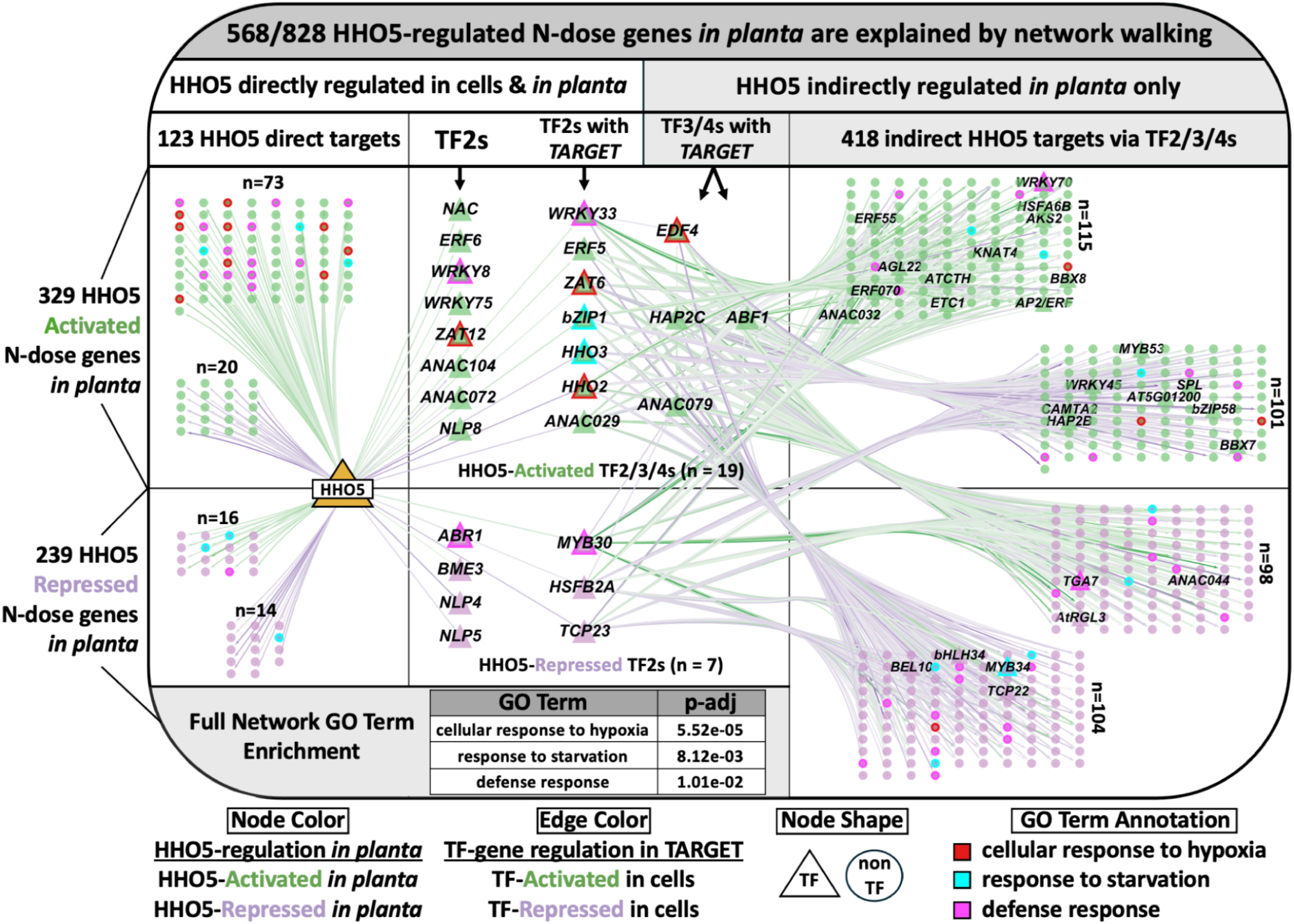
Network walking links the indirect regulatory activity of HHO5 via TF2s to genome wide N-dose responses in planta. Gene Regulatory Network (GRN) of 568 gene nodes. Every node is an N-dose gene that is regulated by HHO5 in planta. Green nodes denote genes activated by HHO5 in planta, and purple nodes show genes repressed by HHO5 in planta. All edges are validated TARGET TF-gene direct regulatory relationships, where green shows activated and purple shows repressed. Top significantly enriched gene ontology terms are shown, and annotated in the network using node border color. TF nodes are shown as triangles, and are labeled by gene name.

The resulting GRN has a bias towards gene activation. For example, of the 123 (non-TF) TARGET nodes downstream of HHO5, 73 of the N-dose genes are activated by HHO5 both in TARGET (green edges) and *in planta* (green nodes) (**Figure 6**). Directionally between TARGET and *in planta* data did not always agree, meaning there may be additional regulatory effects downstream of the TARGET edges in our network. Looking at directionality *in planta* (node color), 329 N-dose genes downstream of HHO5 are activated, while 239 are repressed.

Furthermore, there are 53 total N-dose responsive TFs downstream of HHO5, and 40 of the 53 TFs are HHO5-activated *in planta* (**Supp Figure 10**). Many HHO5 TF2s are well studied regulators of plant N and stress signaling (See Discussion). This network analysis highlights the importance of HHO5 indirect gene activation in controlling N-dose related gene expression via HHO5 TF2s.

HHO5 transcripts are expressed specifically in phloem cells (**Figure 2**), and mutating HHO5 decreases the rate of primary root elongation in response to N-dose (**Figure 3**). To better understand how HHO5 activity in phloem cells may be regulating N-dose dependent primary root growth, we analyzed the cell type specificity of the N-dose genes in the HHO5 network walking GRN. We found 64 network-validated HHO5 target genes to be expressed specifically in phloem cells (**Supp Fig 11A**). Validated HHO5 target genes expressed in phloem cells are enriched for genes related to nitrate import, regulation of root development, and response to starvation (**Supp Fig 11B**). These phloem-specific HHO5 target genes include many well studied N-signal regulating HHO5-TF2s such as *bZIP1, ZAT6, RAV1, HSFB2A,* and *TCP23* (**Supp Fig 12**). Therefore, mutating HHO5 perturbs N-dose and root developmental signals that are expressed in phloem cells, which could explain the N-dose dependent growth defects in *hho5* mutant roots.

## Discussion

Despite sharing conserved *MYB* DNA binding domains and Hydrophobic Globular Domains (HGDs) typical of the HHO TFs family, HHO5 displays distinctive structural differences from other HHO TFs^24^. *HHO1*, *HHO2*, and *HHO3* each contains a single 5’ *ERF-associated Amphiphilic Repressor* (*EAR)*-like domain, which facilitates protein-protein interactions (with EAR domains, and non-EAR-protein domains) and confers target gene repression^25,26^. *HHO5* is unique in carrying two EAR-like domains, located at both the 5’ and 3’ ends. Additionally *HRS1*, *HHO4*, and *HHO6* don’t contain any^24^. The additional repressive EAR-like motif at the 3’ end of *HHO5* may enhance HHO5’s interaction potential with partner TFs, suggesting a possible mechanistic basis for its functional divergence compared to the redundant *NIGT1* clade.

Though *HHO5* is structurally divergent, the HHO family of TFs seem to function largely as direct transcriptional repressors^6^, which is consistent with their EAR-like domains. In this context, direct repression refers to the association between direct contact between TF and DNA and corresponding target gene repression (**Figure 5**). However the association between direct DNA binding by HHO TFs and target gene repression engenders the question; How does HHO5 activate gene expression? In this study we propose that gene activation by HHO5 can be accomplished by two possible (and not mutually exclusive) explanations.

First, we found that HHO5 regulates multiple downstream TFs (TF2s) that are well established gene activators (e.g., *bZIP1, WRKY33, WRKY75*)^17,27,28^. Thus HHO5-TF2s activate target gene expression after N-stimulus via HHO5 (**Figure 5**). We also found instances where HHO5 directly repressed other repressors such as *HHO2* and *HHO3* in TARGET, creating a double-negative effect that ultimately leads to gene activation (**Figure 6**). This TF2 mediated activity can explain why a gene is regulated by HHO5 *in planta*, but not directly regulated by HHO5 in the cell-based TARGET assay. Supporting this hypothesis, our network walking analysis demonstrated that incorporating experimentally validated TF2 activity into the network accounted for almost 70% of the HHO5 regulated N-dose responsive genes observed *in planta*. These findings position HHO5 as a key regulator of N signaling through hierarchical transcriptional control of other important TFs.

The TF2-based hypothesis of HHO5 gene activation however does not explain how HHO5 itself can bind and activate target genes. If direct DNA binding by HHO5 confers target gene repression, then indirect HHO5 DNA binding via partner TFs may explain gene activation directly downstream of HHO5. For example, the HHO5 activated genes (in the TARGET assay, **Figure 5B**) are highly enriched for the *WRKY* family cis motif, and *WRKY8*, *WRKY33, and WRKY75* are N-dose responsive direct target genes of HHO5. Thus *WRKY* TFs may be aiding HHO5 in activating N-dose gene expression.

Our second hypothesis explaining gene activation downstream of *HHO5* thus involves the activity of interacting partner TFs. The presence of the extra EAR-like domain in *HHO5* that facilitates protein-protein interactions complements this hypothesis well. We propose that HHO5 would be important for guiding the TF complex to specific target genes, while the interacting WRKY TF would be responsible for directly contacting the promoter DNA of those activated targets. This hypothesis is further supported by the fact that many WRKY TFs contain EAR domains, meaning WRKY TFs are structurally compatible partners for physical interaction with HHO5^29^. Also, the HHO5 activated genes are highly enriched for immunity related functional terms (**Figure 4**), and the WRKY TFs are well characterized master regulators of plant immunity^30,31^. Further study may reveal that WRKY TFs and HHO5 cooperatively activate gene expression.

Previous research validated HHO5 protein-protein interaction with ULTAPETALA1 (ULT1), which earned *HHO5* the additional name *ULTRAPETALA1-Interacting-Factor-1* (*UIF1*)^7^. *ULT1* is described as an “anti-repressor” in floral meristem regulation which is consistent with our model of combinatorial gene activation by HHO5 and a partner TF. Therefore there are validated HHO5 partner TFs that could aid in HHO5 target gene activation. Another way in which HHO5 is diverged from the *NIGT1* TFs is with its distinct root spatial expression profile.

While the NIGT1 TFs and HHO5 have similar protein domains, HHO5 is expressed in different cell types, meaning HHO5 has access to different partner TFs and chromatin contexts. In summary, the direct repressive DNA binding of HHO TFs is conserved, however the biological context in which the HHO TFs are placed leads to different functional outcomes on plant

N-signaling. Transcriptional dual activity (activating and repressing) is of great interest in the genomics field, and HHO5 may be an interesting candidate to study how dual activity may mechanistically occur^21^.

The specific expression of HHO5 in phloem cells may explain why perturbing HHO5 has a uniquely strong effect on N-dose signals. The more redundant NIGT1 TFs are expressed broadly in many root cell types, meaning they likely functionally compensate for each other.

NIGT1 TF redundancy is apparent in the fact that additive mutants are needed to achieve a N-related plant phenotype effect^6^. HHO5 on the other hand has specific expression in phloem, and single mutation of HHO5 is sufficient to strongly dampen plant N-dose signaling. Phloem cells are important for the transporting of N in the form of nitrogenous amino acids^32^. We found that HHO5 regulates nutrient starvation and nitrate import specifically in phloem cells (**Supp Fig 11**). This localization to phloem cells may contribute to the N-dose dependency of HHO5 regulation, which we quantified as a significant genotype*N-dose interaction effect on gene expression and phenotype outcomes in response to HHO5 perturbation *in planta.* The

phloem-localization of some HHO5 and N-dose regulated genes could explain the decreases in plant fitness (seed production) and seed N content detected in *hho5* mutants. Thus, we propose that HHO5 is an important regulator of systemic plant N signaling both at the gene expression and phenotype levels.

In conclusion, HHO5 is unique among the HHO family of TFs, in that it is the only N-dose responsive HHO TF that is phylogenetically diverged from the four redundant *NIGT1* TFs (**Figure 2**). This combined with our findings that *HHO5* responds to N-signals within minutes^3^, and is specifically expressed in phloem cells (**Figure 2**), made *HHO5* a strong candidate regulator of plant N-signaling. Our findings reveal that HHO5 regulates the N-dose response of over 800 genes in Arabidopsis roots. Similarly to the NIGT1 TFs, HHO5 direct DNA binding leads to repression of target gene expression. This means gene activation downstream of HHO5 is associated with indirect DNA binding, or indirect regulatory effects by downstream

HHO5-TF2s. We validated the indirect effects of HHO5 on N-dose responses using a network walking approach which linked the activity of HHO5-TF2s to regulation of over 400 N-dose genes *in planta*. Opposing the canonical repressive roles of HHO TFs, we found that mutating HHO5 actually dampened N-dose responses at the phenotypic level. The novelty that HHO5 activates phenotypic responses to N-dose highlights the importance of studying both the direct and indirect regulatory effects of the phylogenetically diverged HHO TF, HHO5, at a systems biology level.

## Materials and Methods

### HHO5 mutants

Mutants *hho5.1* (SAIL_806_F06) and *hho5*.2 (SALK_077802) hosted T-DNA insertions in the first and fifth exons respectively (**Supp Fig 13A**). Homozygous lines of each allele were obtained and their genotypes confirmed by PCR after two backcrosses to the parental Col-0 ecotype to remove extra T-DNA insertions. Left border sequencing of the T-DNA was conducted to verify the insertion site of each allele. To test for *hho5* transcript, total RNA was extracted from 2-week-old wild-type and mutant plants grown on Murashige and Skoog (MS) medium and reverse transcribed. Both, *hho5*.1 and *hho5*.2 showed the absence of *hho5* transcript in seedlings when using primers designed to amplify 3’ UTR transcript (**Supp Fig 13B** and **Table S8**).

### HHO5 mutant phenotyping

Seeds were sown on vertical 120-by-120 mm square plates containing 50 mL of MS basal salts, supplemented with 3 mM sucrose, and 0.8% (w/v) agar at pH 5.7 and KNO_3_ or NH_4_NO_3_. Surface-sterilized seeds were aligned on the plate surface; mutant lines and wild-type Col-0 were placed on the same plate. Primary root length was scored every 3 days. The root length was quantified using ImageJ 1.52a software (National Institutes of Health). At the end of the experiment, plant shoots were extracted and fresh and dry weight were measured. For the measurement of dry weight, tissue was dried at 70°C for 3 days, and weighed to determine dry weight. Silique length was evaluated in mature Col-0 and *hho5* mutant plants. 10 plants of each line were used for silique and seed counting (3 siliques per plant).

Siliques from the middle part of the main inflorescence were analyzed. Seed number per silique was recorded for each line. All data are expressed as means ± SEM. We found that phenotypic responses were the same regardless whether the N source was present as KNO_3_ or NH_4_NO_3_ (**Supp Fig 5**). Comparisons among groups were shown using two-way analysis of variance (ANOVA) with Bonferroni post-tests. Interaction effects (genotype*N-dose) were tested using the aov() function in R. Total N content in seeds was determined by the Kjeldahl method.

#### Histochemical staining and image analysis

The histochemical analysis of GUS reporter enzyme activity was performed as described by Weigel.^33^ The staining patterns of GUS in the roots were visualized under the Olympus BX51 microscope (Olympus, Tokyo, Japan) with differential interference contrast optics. p*1kbHHO5::GUS* seedlings were grown for 10 days on MS media, supplemented with 1 mM KNO_3_ and 1 % sucrose. Plants were grown under long-day conditions (16 hours light, 8 hours dark), with a light intensity of 120 μmol m^-2^s^-1^ at constant temperature of 22 ⁰C. After 10 days, plants were starved of N for an additional day. 2 hours after subjective dawn on the 12^th^ day, plants were treated with no N, 5 mM or 10 mM KNO_3_ for 2 hours (**Figure 2C**).

#### HHO5 N-dose course

We grew ∼100 seedlings Col-0, *hho5.1* and *hho5.2 Arabidopsis* in hydroponics. Seedlings were grown for 13 days on MS media, supplemented with 1 mM KNO_3_ and 1 % sucrose. Plants were grown under long-day conditions (16 hours light, 8 hours dark), with a light intensity of 120 μmol m^-2^s^-1^ at constant temperature of 22 ⁰C. After 13 days, plants were starved of N for an additional day. 2 hours after subjective dawn on the 15^h^ day, plants were treated with 1 of 6 N doses for 2 hours. Doses were 0, 0.002, 0.02, 0.2, 2 and 20 mM KNO_3_ + NH_4_NO_3_. Additionally, 1 set of plants were treated with 20 mM KCl to control for the effect of potassium. Plants were treated in triplicate, resulting in 63 samples total. After a 2 hour incubation period, root tissue was excised and flash frozen in liquid nitrogen. RNA was extracted from whole roots using the Qiagen RNeasy kit with on-column DNAse digestion, and quality checked using Agilent Tapestation HSRNA Kit. mRNA was isolated using oligo-dT purification from 1 μg of total RNA (Thermo Fisher Scientific) and libraries made using NEBNext RNA Kit. Libraries were sequenced using Illumina NextSeq with 1x75 bp read chemistry.

Resulting reads were aligned to the TAIR10 genome using Tophat^34^, and aligned to the Araport 11 *Arabidopsis* annotation^35^ using HT-Seq^36^ to call gene counts.(Raw Gene Expression can be found in Supp Table S9).

### Calling and visualizing N-dose DEGs with DESeq2

All Code is available in the supplemental RMD. Raw gene counts were used in DESeq2^37^ for normalization. VST normalized counts of the top 5000 most variable genes were put into the DESeq2 PlotPCA function to visualize the entire transcriptome dataset (**Supp Fig 1**). N-dose DEGs were called using the DESeq2 LRT Test full model ∼ Genotype + Dose + Treatment, where genotype referred to WT, *hho5.1*, and *hho5.2* mutants. Dose referred to the numerical dose of nutrient treated. The Treatment term differentiated the N treatment from the KCl treatment control. 0 mM was classified as part of the control treatment. The LRT test reduced model ∼ Genotype + Treatment was used to call 4,644 N-dose DEGs (p.adj <0.01, **Figure 1A**). DEGs were clustered and visualized using the DEGReport package in R using consensusCluster set to True and minimum cluster (minc) set to 10 genes^38^. (**Figure 1A**). Gene Ontology enrichment analysis was completed on these clusters using the clusterProfiler package in R, with ontology set to “BP”, FDR and p-value set to 0.05, and KetType set to TAIR **Figure 1B**)^39^. Additional clusters were identified using DEGReport with consensusCluster set to False (**Supp Fig 2**). N metabolism genes were identified using the KEGG database.^14^ These N-metabolism genes were intersected with the 4,464 N-dose genes, and scaled gene expression fo the N-dose responsive N-metabolism genes was visualized in a heatmap using the ComplexHeatmap package in R^40^. The expression of individual genes was plotted using the DESeq2 PlotCounts function and ggplot2^41^.

### Analysis of HHO Family N-Dose response

Gene expression of the HHO TFs were plotted across N-doses using ggplot2 (**Figure 2A**). Protein sequences of the seven HHO TFs were acquired from TAIR.org and used as input for ClustalOmega to generate a protein phylogeny tree^42^. Cell-Type expression profiles were generated by plotting a heatmap of the average gene expression of the HHO TFs across cell types from the Arabidopsis root cell type atlas from Li et al 2016 (**Figure 2B**)^15^.

### Calling HHO5-regulated N-Dose genes in planta with DESeq2

Using the same DESeq2 object as before, HHO5 genotype DEGs were called using the standard DESeq2 wald test for each allele separately. This was accomplished by setting contrasts for (*hho5.1* vs WT) and (*hho5.2* vs WT) with p.adj <0.05. We then took the intersection of the *hho5.1* and *hho5.2* DEGs, resulting in 2,894 high confidence DEGs called in both mutant alleles (**Figure 3A**). The significant overlap between these two lists was tested using a hypergeometric distribution test in R (phyper). These 2,894 HHO5-regulated genes were intersected with the N-dose genes, and the 828 genes overlapping represented a significant intersection based on the hypergeometric test (**Figure 3A**). The same clustering packages and parameters were used to visualize these genes as in (**Figure 1A)**. An ANOVA was performed within each cluster to test for significant genotype*interaction effects using the aov() function in R.

### Analysis of HHO5 direct binding and gene regulation

HHO5 TARGET RNA-seq data was acquired from ConnecTF.org (**Figure 5A**)^23^. Gene ontology analyses on these lists were performed as described for **Figure 1B**. HHO5 DAP-seq and amp-DAP-seq datasets were generated by O’malley and Huang et al 2015^19^. Reads were aligned to the TAIR10 genome, extended to the mean sequencing fragment size (200 BP), and normalized using an RPKM normalization. This resulted in bigwig files that were used as input into the R package EnrichedHeatmap to generate spatially resolved DAP-seq binding plots (**Figure 5B**)^43^. Significance of the intersection between HHO5 binding and regulation was completed using the hypergeometric test in R. Cis-motif images were acquired from the Ecker lab plant cistrome database. Cis-motif enrichment analysis was completed in ConnecTF.org using the 500 BP promoters of each HHO target gene^23^.

### Network Walking

The gene regulatory network (GRN) was generated using the network functions in ConnecTF. The 828 HHO5-regulated N-dose DEGs were used as input target genes, and edges were restricted to TARGET data (direct gene regulation) only for TFs that were both N-dose regulated and downstream of HHO5 in cells and *in planta*. TFs downstream of HHO5 were then mapped to other N-dose TFs that were regulated by *HHO5* in planta, but not in cells (indirectly regulated). The network was visualized using cytoscape^44^.

## Acknowledgements/Funding

W.E.H was supported by NIH award R35GM138143 and QBIST NIH T32 Training Program of NYU biology grant 1T32GM132037-01. J.P.M and M.O are investigators of the National Research Council (CONICET) from Argentina. This work was supported by ANPCyT (PICT 2015-3052 and PICT 2019-4328) and PIP 11420100100242 by CONICET. F.R was supported by the CONICET-PUE grant. This work was supported by NIH Grant RO1-GM121753 to GMC. We acknowledge the Zegar Family Foundation for their generous support. This work was supported in part through the NYU IT High Performance Computing resources, services, and staff expertise. We thank NYU Gencore for their generous support and resources.

## Author Contributions

J.S., M.O. and G.M.C. conceived and designed the experiments; W.E.H. and J.S. conducted the experiments, analyzed the sequencing data and performed the bioinformatics and statistical analyses. M.O, F.R and J.P.M. performed physiological experiments. W.E.H., J.S., S.S.C.H., J.P.M., G.M.C. and M.O. and interpreted the results. W.E.H., J.S., G.M.C. and M.O. wrote the article with contributions from all authors. All authors read and approved the final manuscript

## Accessions and Data Availability

Both *hho5* T-DNA mutants are available for ordering from the Arabidopsis Biological Resource Center (ABRC) under the following accessions: *hho5.1* (SAIL_806_F06), *hho5.2* (SALK_077802). The *in planta* transcriptome dataset from this study (**Supp Fig 1**) was made available at the Gene Expression Omnibus (GEO) database under accession number PRJNA512225. Gene expression counts and associated code were made available as **Table S9** and **Supp File 1**, respectively.

## Supplemental Figures

**Supplemental Figure 1:**
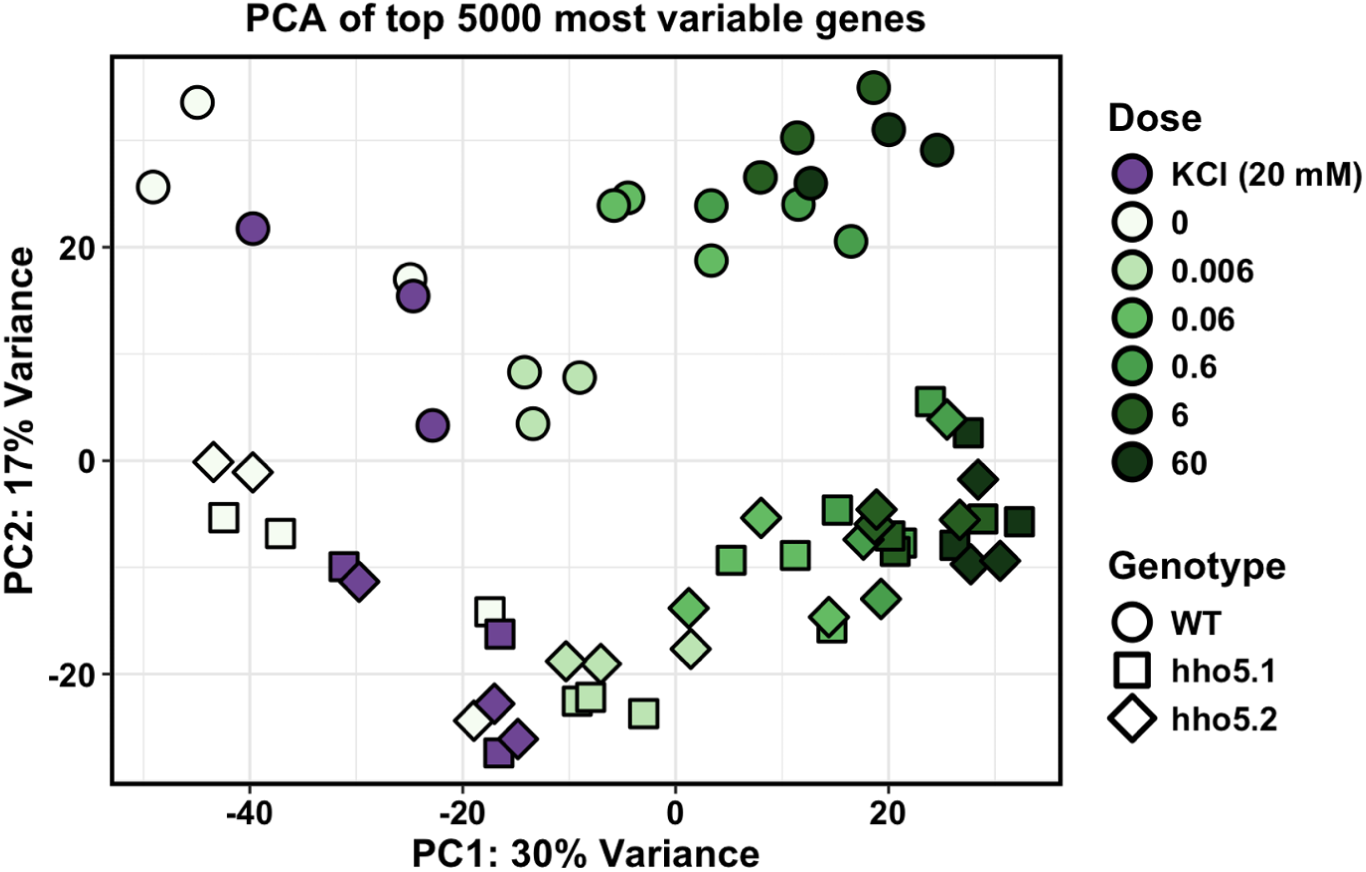
Transcriptome-wide N-dose responses in Arabidopsis roots. Principal Component Analysis (PCA) of the top 5000 most variable genes in Arabidopsis roots, showing variation across genotypes (shape) and nutrient treatments (colors).

**Supplemental Figure 2:**
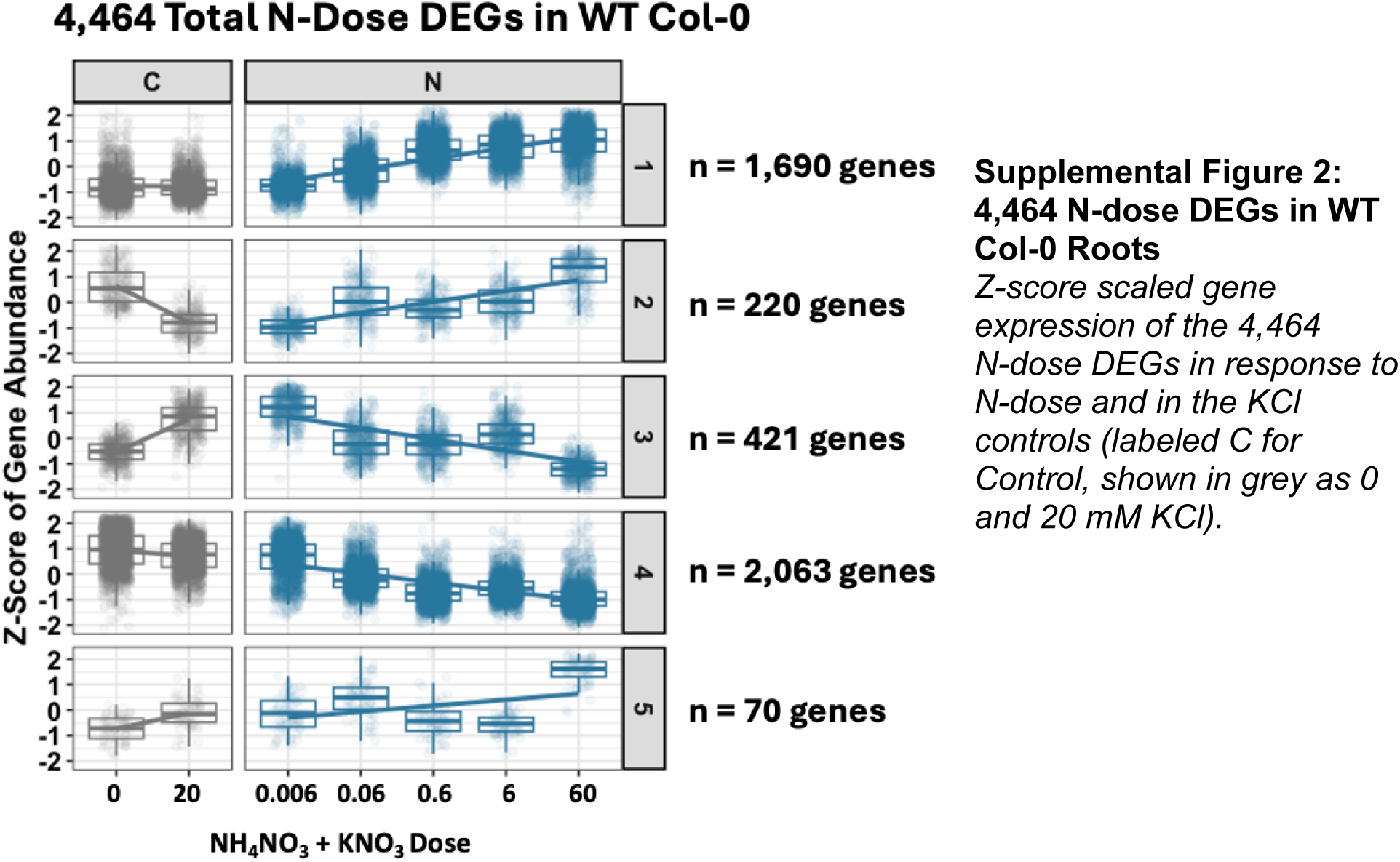
4,464 N-dose DEGs in WT Col-0 Roots. Z-score scaled gene expression of the 4,464 N-dose DEGs in response to N-dose and in the KCl controls (labeled C for Control, shown in grey as 0 and 20 mM KCl).

**Supplemental Figure 3:**
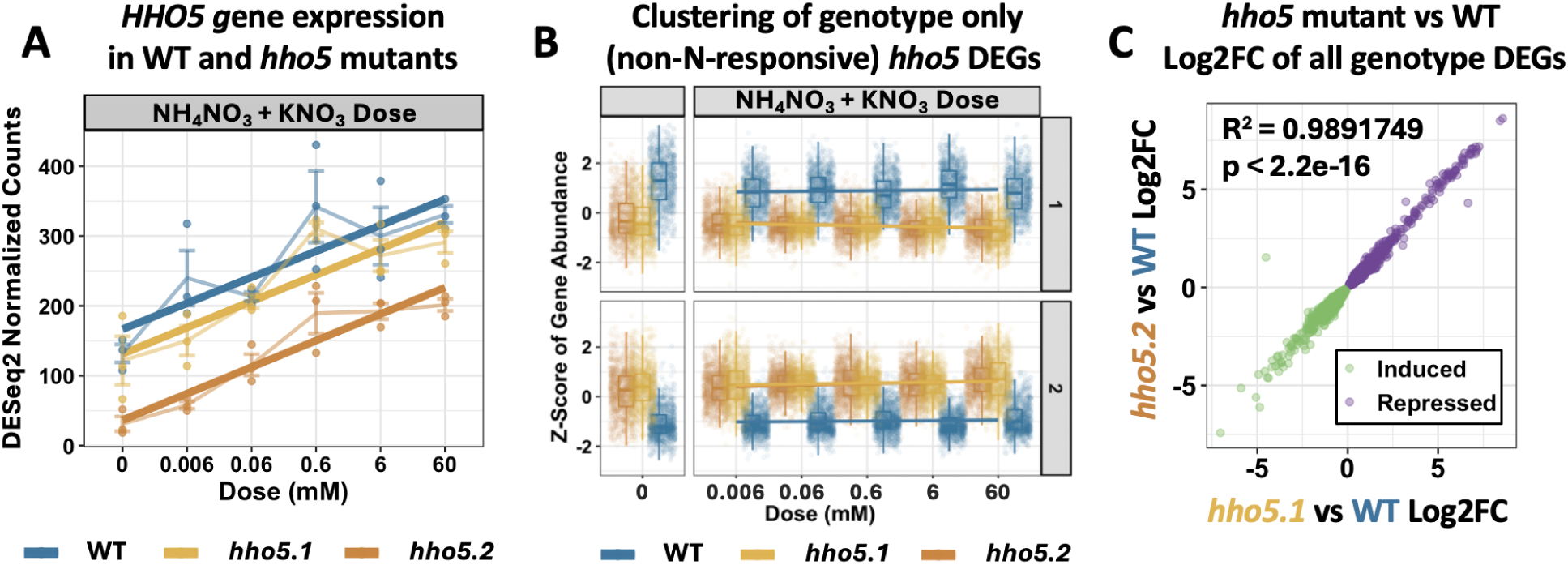
HHO5 regulates thousands of genes in planta. A) HHO5 gene expression across genotypes and N-doses. B) Expression profiles of genes activated (top) and repressed (bottom) by HHO5, but not regulated by N-dose. C) Correlation of Log2FC between hho5.1 and hho5.2 mutants for the 2,894 genes regulated by HHO5 in planta.

**Supplemental Figure 4:**
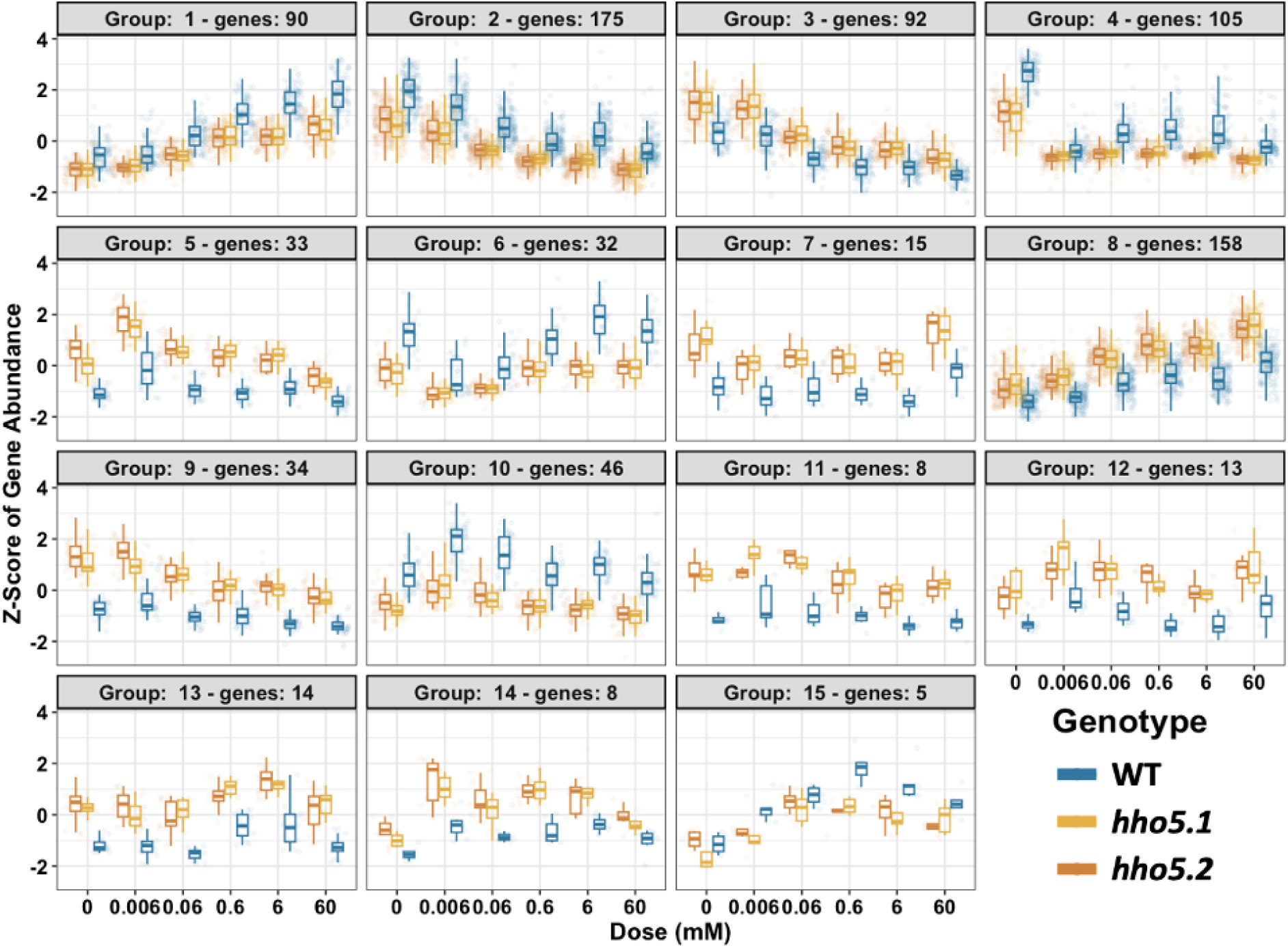
Clustering of 828 HHO5-Regulated N-dose genes. A) 15 gene expression clusters of z-score scaled gene expression across N-doses in WT and two hho5 T-DNA mutants. N-dose represents Total N Dose as described in Fig 1.

**Supplemental Figure 5:**
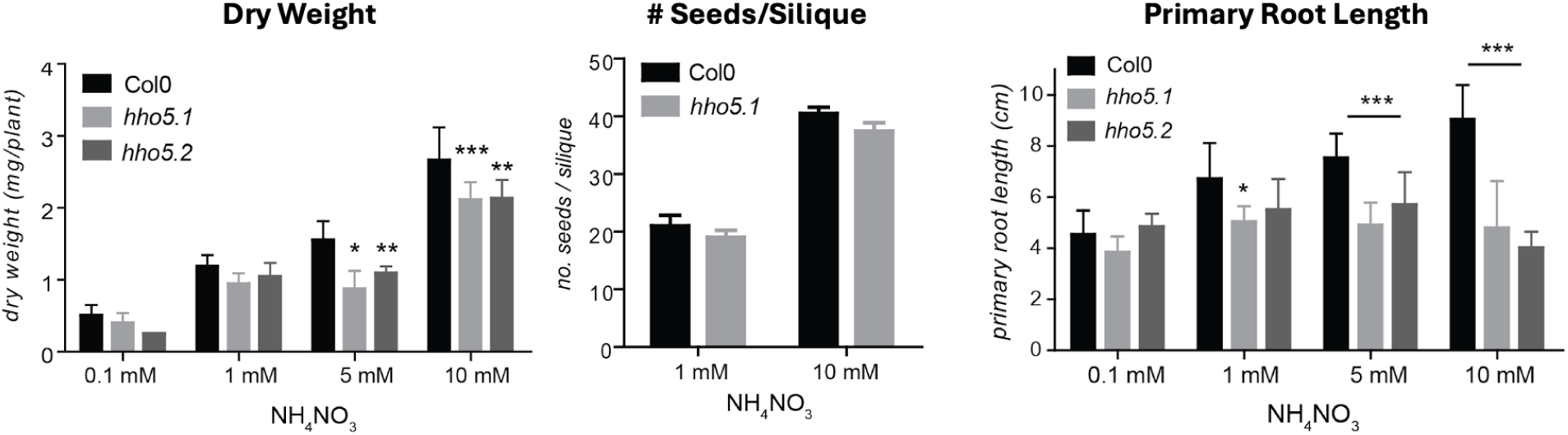
hho5 mutant growth in ammonium nitrate treatments. A) Dry weight, B) # Seeds per silique, and C) Primary Root Length in WT and two hho5 null T-DNA mutants in response to different doses of ammonium nitrate.

**Supplemental Figure 6:**
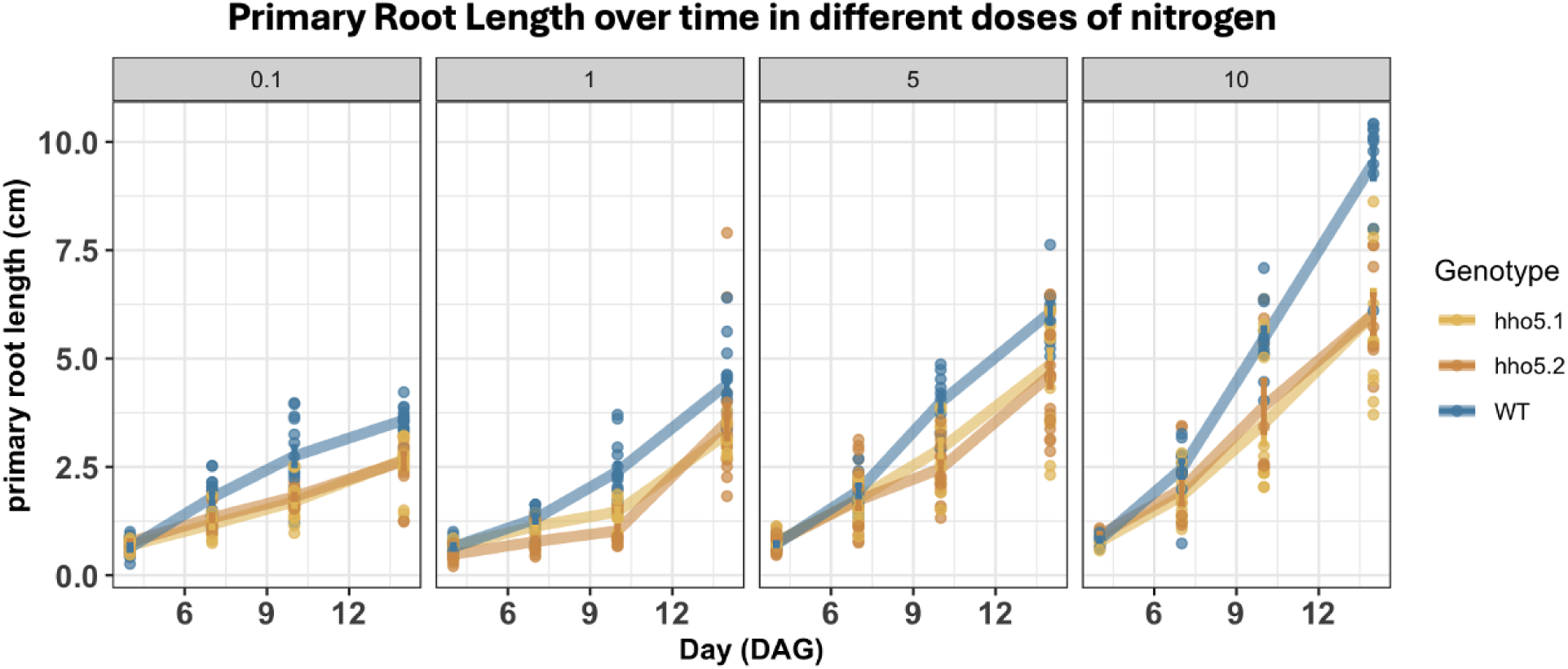
hho5 mutant primary root length responses to KNO_3_ treatment. Plots of primary root lengths across 4 doses of KNO_3_ (0.1, 1, 5, 10 mM) across 4 time points.

**Supplemental Figure 7:**
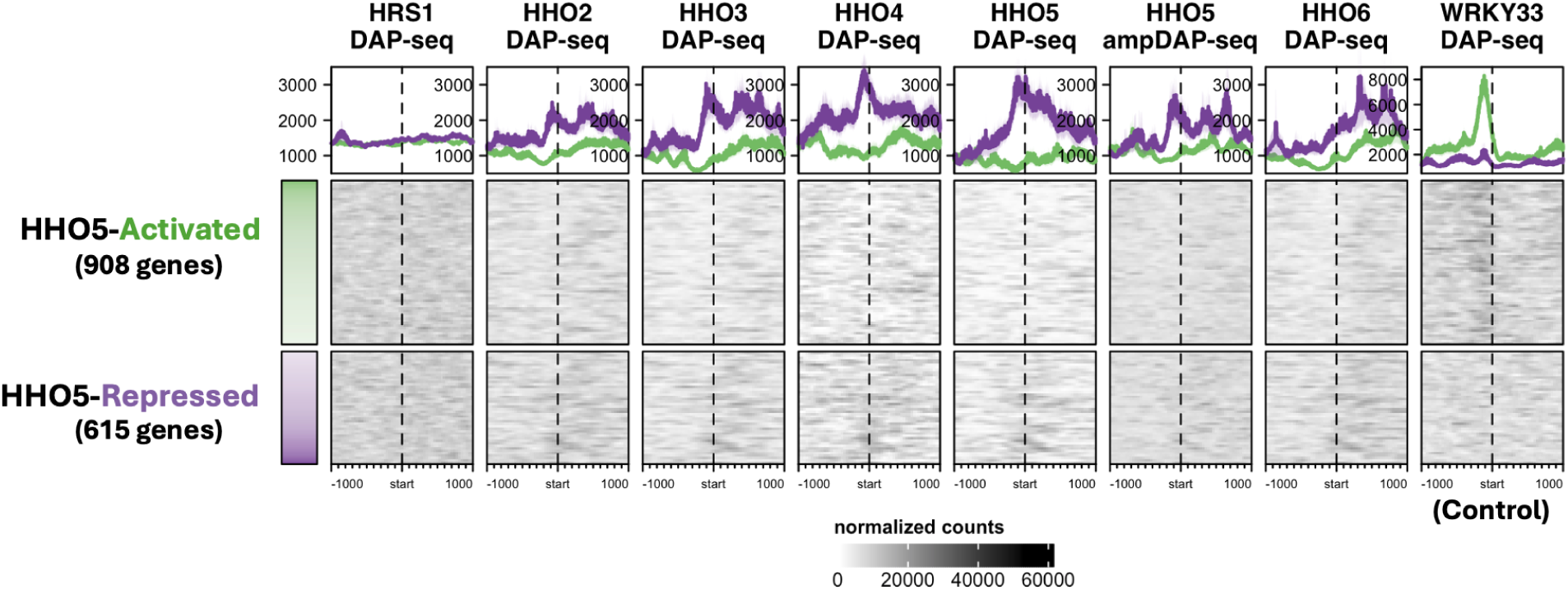
The HHO family preferentially binds the HHO5-repressed genes. Analysis of HHO5 DAP-seq and ampDAP-seq binding^19^ over the HHO5 directly regulated genes from TARGET assay^18^. This is plotted +/- 1 kb from the gene transcription start sites. DEGs are ordered by Log2FC. Lack of binding in HRS1 is likely due to its very low number of peaks (883), and a low fraction of reads under the peaks (∼3%).

**Supplemental Figure 8:**
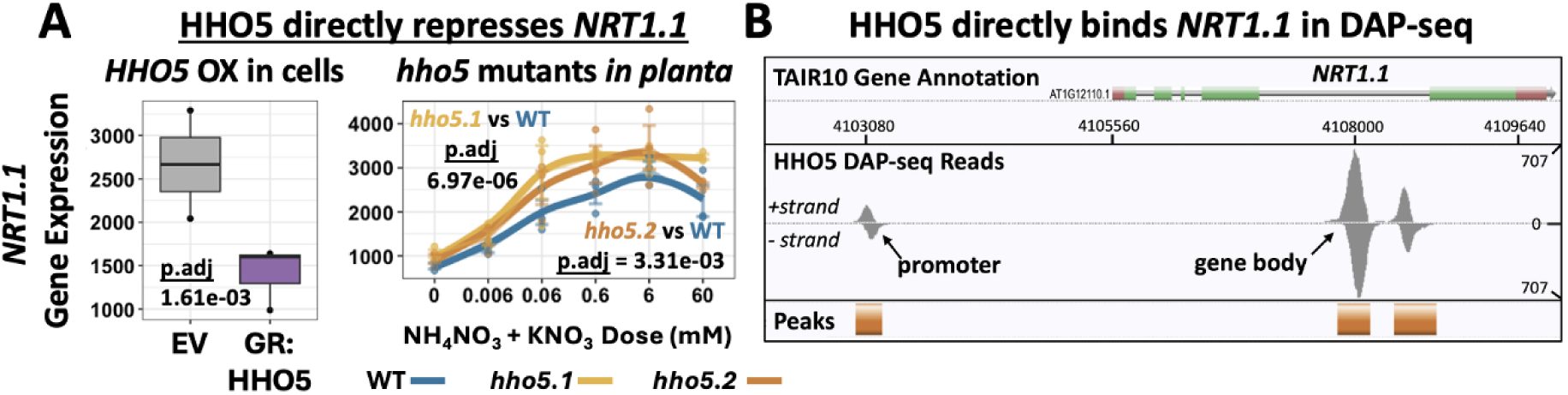
HHO5 directly binds and represses NRT1.1. A) Gene expression of NRT1.1 (AT1G12110) in the TARGET assay empty vector (EV, no TF control) and the GR:HHO5 over-expressing cells. Plotted next to the gene expression of NRT1.1 in WT and hho5 T-DNA mutants across N-doses from this study. B) DAP-seq DNA binding of HHO5 over NRT1.1 in the Ecker lab cistrome genome browser^19^ .

**Supplemental Figure 9:**
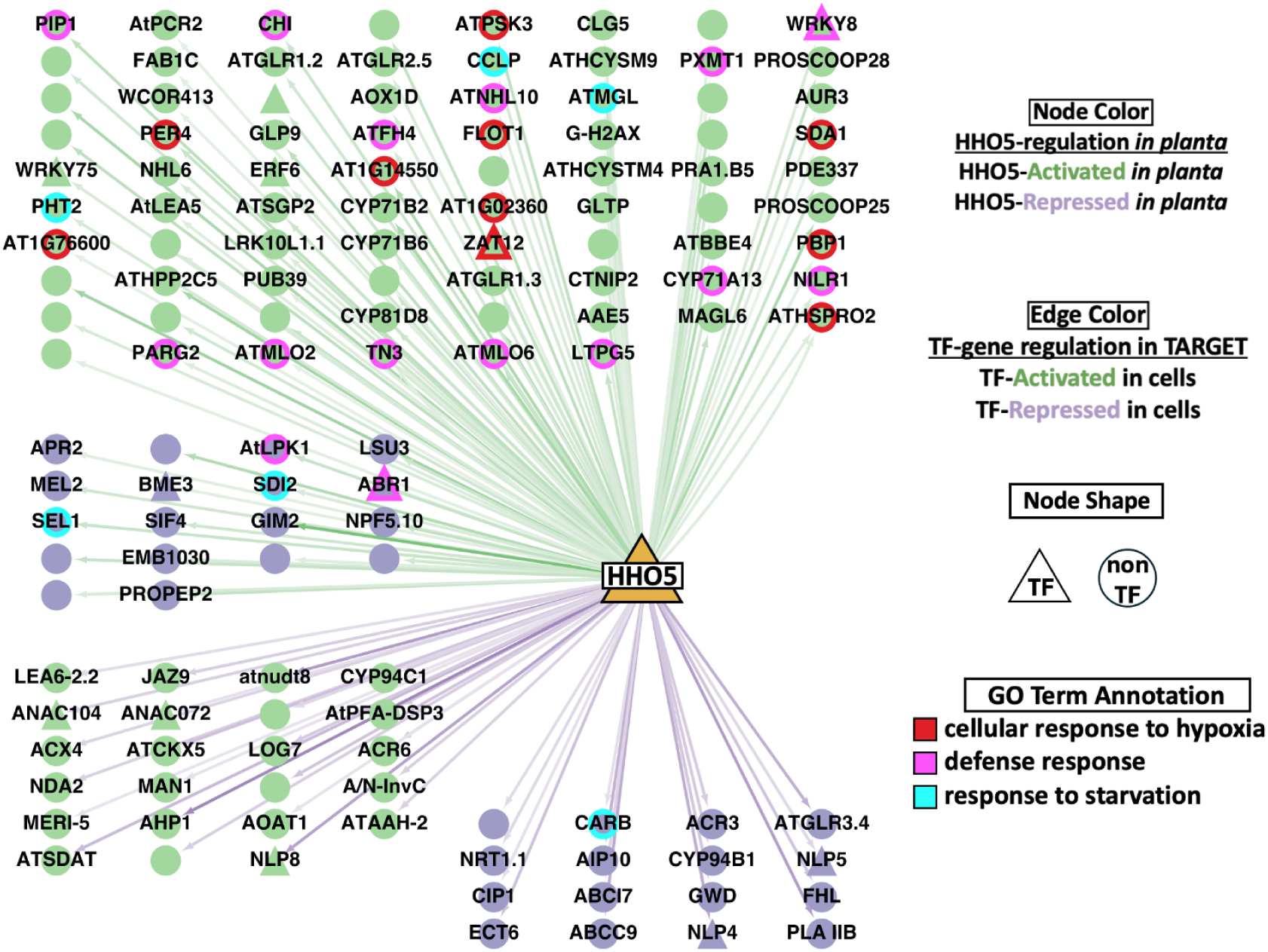
135 Direct Targets of HHO5 in cells and in planta. A subset of the network walking GRN highlights 135 N-dose genes downstream of HHO5, most of which (101/135) are directly activated by HHO5 (in the TARGET system, green edges).

**Supplemental Figure 10:**
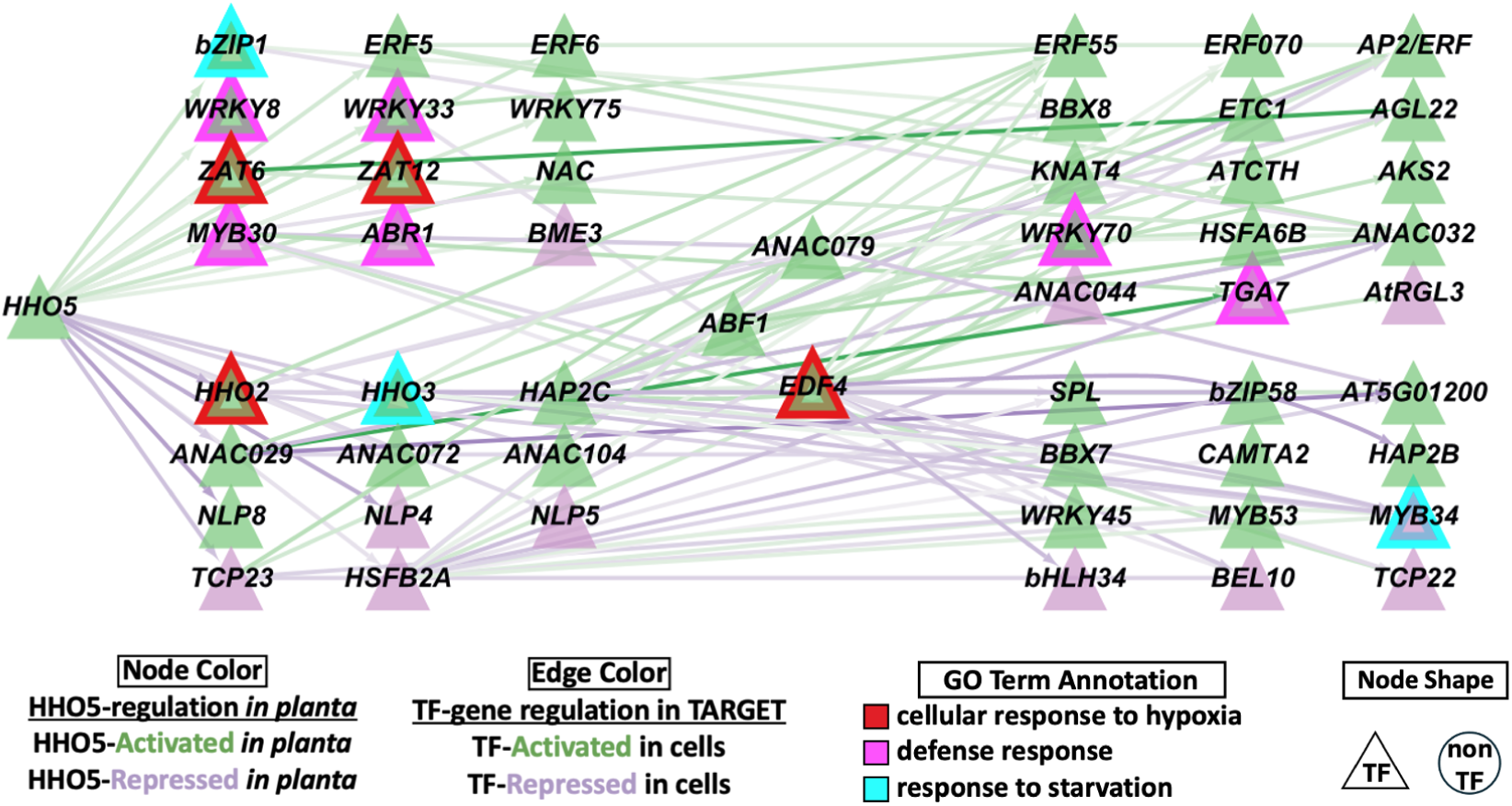
53 N-dose TFs downstream of HHO5 tend to be HHO5-activated. Network showing all regulatory edges between HHO5 and the 53 validated N-dose responsive TFs downstream of HHO5 using TARGET data. 40 of the 53 are activated by HHO5 in planta. Line color denotes intensity of Log2FC in TARGET.

**Supplemental Figure 11:**
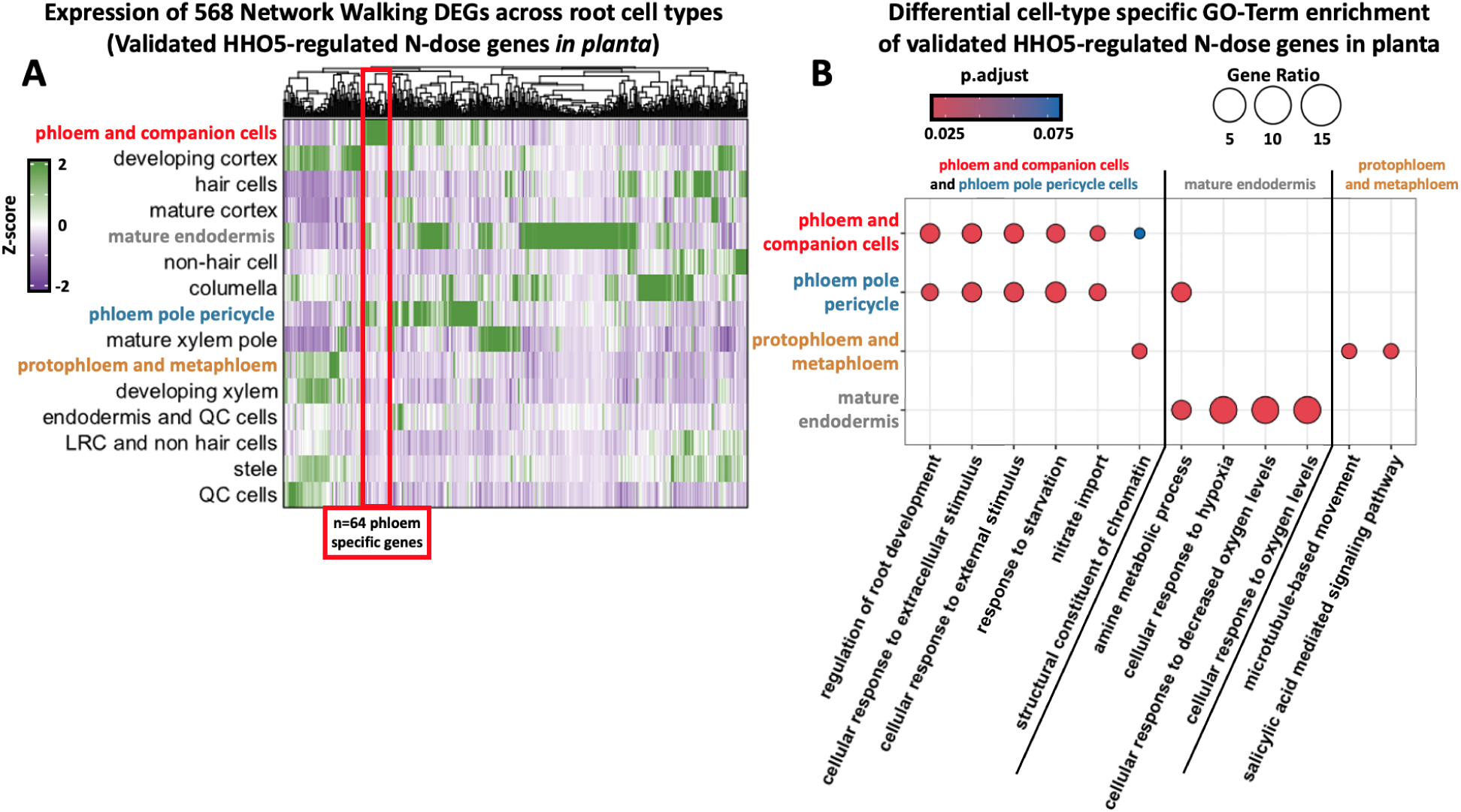
The phloem-expressed HHO5-regulated N-dose genes validated by Network Walking are enriched in genes related to “regulation of root development”. A) Heatmap of 568 validated HHO5-regulated N-dose genes (network walking genes from Figure 5) across the root cell type atlas^15^. Genes highly expressed in phloem and companion cells (Z-score >1) were selected for gene ontology enrichment analysis. B) GO-Term enrichment across cell types, with mature endodermis as a control.

**Supplemental Figure 12:**
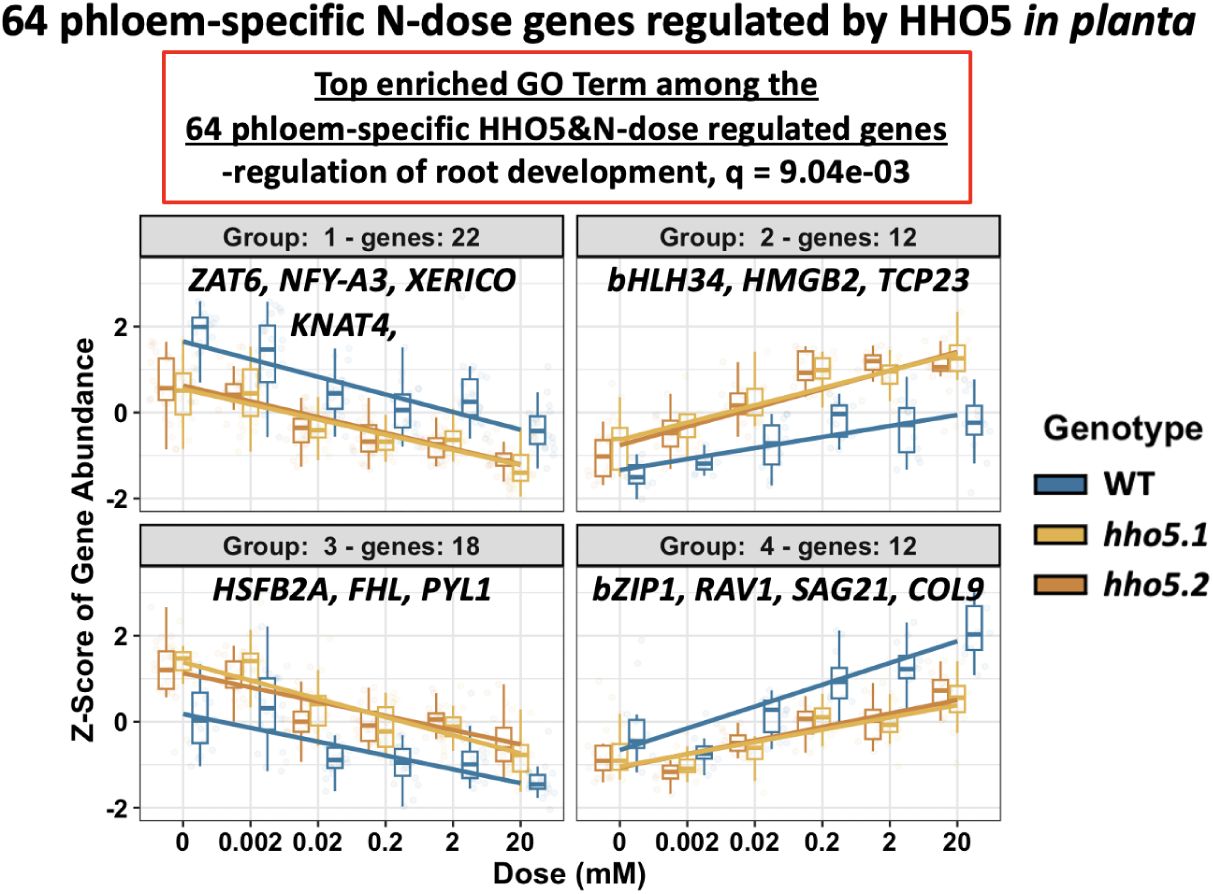
The phloem-expressed HHO5-regulated N-dose genes validated by Network Walking are enriched in genes related to “regulation of root development”. 4 clusters of gene expression profiles from the 64 phloem expressed HHO5-regulated N-dose DEGs (See Supp Fig 11). These 64 genes are enriched for “regulation of root development.”

**Supplemental Figure 13:**
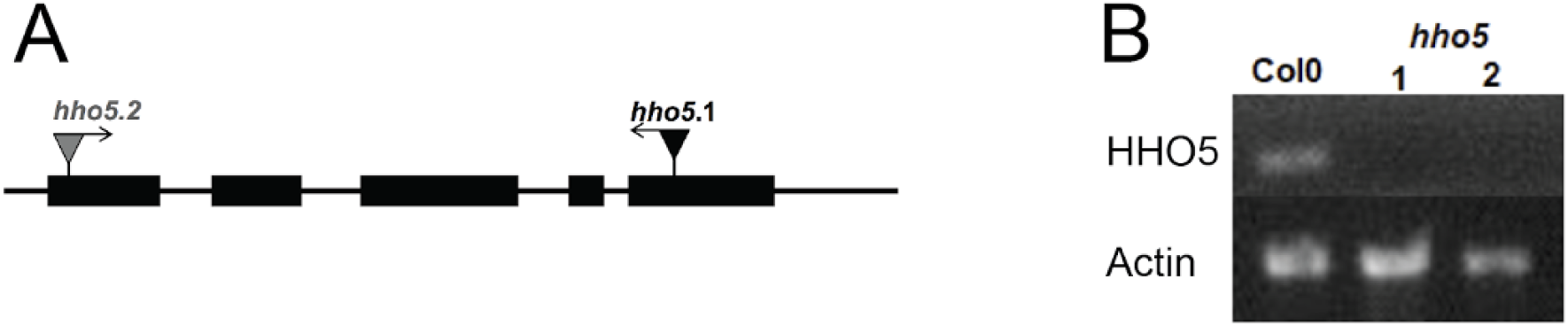
Genotype analysis of two hho5 T-DNA insertion alleles. A) The hho5.1 T-DNA is in the final exon of the gene, while the hho5.2 T-DNA is near the 5’ UTR. B) RT-PCR revealed these are both null mutants, as no amplification of HHO5 was seen in either, compared to WT Col-0 plants as a control. ACTIN was used as a loading control.

